# Systematic mapping and modeling of 3D enhancer-promoter interactions in early mouse embryonic lineages reveal regulatory principles that determine the levels and cell-type specificity of gene expression

**DOI:** 10.1101/2023.07.19.549714

**Authors:** Dylan Murphy, Eralda Salataj, Dafne Campigli Di Giammartino, Javier Rodriguez-Hernaez, Andreas Kloetgen, Vidur Garg, Erin Char, Christopher M. Uyehara, Ly-sha Ee, UkJin Lee, Matthias Stadtfeld, Anna-Katerina Hadjantonakis, Aristotelis Tsirigos, Alexander Polyzos, Effie Apostolou

## Abstract

Mammalian embryogenesis commences with two pivotal and binary cell fate decisions that give rise to three essential lineages, the trophectoderm (TE), the epiblast (EPI) and the primitive endoderm (PrE). Although key signaling pathways and transcription factors that control these early embryonic decisions have been identified, the non-coding regulatory elements via which transcriptional regulators enact these fates remain understudied. To address this gap, we have characterized, at a genome-wide scale, enhancer activity and 3D connectivity in embryo-derived stem cell lines that represent each of the early developmental fates. We observed extensive enhancer remodeling and fine-scale 3D chromatin rewiring among the three lineages, which strongly associate with transcriptional changes, although there are distinct groups of genes that are irresponsive to topological changes. In each lineage, a high degree of connectivity or “hubness” positively correlates with levels of gene expression and enriches for cell-type specific and essential genes. Genes within 3D hubs also show a significantly stronger probability of coregulation across lineages, compared to genes in linear proximity or within the same contact domains. By incorporating 3D chromatin features, we build a novel predictive model for transcriptional regulation (3D-HiChAT), which outperformed models that use only 1D promoter or proximal variables in predicting levels and cell-type specificity of gene expression. Using 3D-HiChAT, we performed genome-wide *in silico* perturbations to nominate candidate functional enhancers and hubs in each cell lineage, and with CRISPRi experiments we validated several novel enhancers that control expression of one or more genes in their respective lineages. Our study comprehensively identifies 3D regulatory hubs associated with the earliest mammalian lineages and describes their relationship to gene expression and cell identity, providing a framework to understand lineage-specific transcriptional behaviors.

**HIGHLIGHTS:**

- Cell lines representing early embryonic lineages undergo drastic enhancer remodeling and fine-scale 3D chromatin reorganization
- Highly interacting 3D hubs strongly enrich for highly expressed, cell-type specific and essential genes
- 3D chromatin features greatly improve prediction of cell-type specific gene expression compared to 1D promoter features
- *In silico* and experimental perturbations identify novel enhancers regulating the expression of two or more genes in early embryonic lineages

## INTRODUCTION

Mammalian development starts with two critical cell fate decisions that give rise to the progenitors of all embryonic and extraembryonic tissues required for proper embryogenesis^1–4^. During the first decision, cells of the totipotent morula segregate into either the inner cell mass (ICM) or the trophectoderm (TE) cells, a polarized epithelial cell layer that gives rise to trophoblast tissues of the placenta. At a later stage, the ICM will generate the pluripotent epiblast (EPI) and the primitive endoderm (PrE) cells which will eventually form the embryo proper and the extraembryonic yolk sac tissue, respectively^5^. Both *in vivo* and *in vitro* studies have uncovered cellular and molecular hallmarks of these early embryonic decisions, including the key signaling pathways (such as Notch, Wnt/B-catenin, Hippo etc.) and DNA-binding transcription factors (TF) that drive lineage specification and segregation^6–8^. However, little is known so far about the downstream non-coding DNA elements and regulatory networks that enforce these early embryonic fates.

Enhancers are essential regulatory elements that-together with TFs-regulate transcriptional activity of gene promoters often over large distances, establishing cell type-specific gene expression programs and hence cellular identities^9, 10^. Chromatin profiling assays, such as ATAC-seq for chromatin accessibility or ChIP-seq for characteristic histone marks (e.g. H3K27ac) have been extremely useful for annotating hundreds of thousands of putative enhancers on a genome-wide scale in various tissues and cell lines^11–15^. However, these assays have limited capacity to assign enhancers to the correct target gene, and to predict their relative regulatory impact on gene expression and cell identity, as shown by reporter assays^16–18^ and genetic or epigenetic engineering^19, 20^. The emergence of 3D chromatin organization as an important regulatory layer of gene expression and cell identity, as other groups and we have shown^21–27^, highlights the necessity of studying enhancer specificity and activity in the context of their 3D neighborhood. This includes the specific long-range interactions of a given enhancer with one or more target genes, the insulating boundaries that may restrict enhancer function and the larger-scale compartmental organization^28–35^. Indeed, genome-wide Chromosome Conformation Capture (3C)-based chromatin assays, such as Hi-C^36^, Capture-C^37, 38^, Micro-C^39, 40^ or HiChIP^41–45^ in various cellular contexts have enabled mapping of 3D enhancer-promoter interactions that are both highly complex and largely cell-type specific. These 3D networks have significantly improved enhancer-promoter assignments and predictions of enhancer functionality compared to traditional approaches based on linear proximity^10, 46–48^.

So far, construction and analysis of 3D networks has not been utilized to dissect and predict regulatory principles that govern early cell fate decisions. Applying genomics technologies to study early embryogenesis *in vivo* is particularly challenging due to the limited cell numbers in the mouse preimplantation blastocyst. Although recent advanced technologies enabled mapping of the transcriptional programs, chromatin states and large-scale chromatin organization of single-cells in various early embryonic stages, they often suffer from poor genomic resolution^49–54^. On the other hand, embryo-derived stem cell lines, known as Trophoblast Stem Cells (TSC), Embryonic Stem Cells (ESCs) and eXtraEmbryonic ENdoderm cells (XEN) have been valuable tools for studying mechanisms that govern the early embryonic lineages of TE, EPI and PrE derivatives, respectively^55–59^. Among them, mouse ESCs that represent the naive EPI state have been extensively characterized by us and others using multiple-omics assays and functional screens ^26, 27, 60, 61^. However, only a few recent studies have started to shed light on the enhancer landscape and 3D chromatin organization of TSC and less so of XEN cells^62–69^ whilst direct comparisons of the 3 lineages are missing.

In this study, we performed multi-omics analysis to comprehensively map the 1D enhancer landscapes and 3D putative regulatory interactions in ESC, TSC and XEN cells as a means of identifying *cis*-regulatory elements and 3D networks that govern early embryonic lineages. Our integrative analysis revealed an extensive enhancer remodeling and 3D rewiring among these closely related lineages and uncover specific links to their transcriptional programs. By applying a Random Forest machine learning approach using various 1D and/or 3D features, we determined important 3D variables that enable better prediction of transcriptional behaviors, such as levels and cell-type specificity of gene expression or gene coregulation. Using an optimized 3D predictive model, which we coin 3D-HiChAT, we also performed genome-wide *in silico* perturbations to predict putative enhancers with regulatory impact on one or more genes in each lineage. Finally, with a series of experimental perturbations in ESCs and XEN, we identified several novel functional enhancers and 3D hubs that control expression levels of one or more developmentally-relevant genes, including *Tfcp2l1 and Klf2 in ESC* and *Mycn* or *Lmna* in XEN cells^70–72^. In conclusion, our study provides a high-resolution 3D atlas of candidate regulatory interactions in early mouse embryonic lineages and reveals novel regulatory principles that determine the levels and cell-type specificity of gene expression.

## RESULTS

### Early developmental decisions are accompanied by drastic enhancer remodeling linked to lineage-specific transcriptional programs

To model and characterize the chromatin regulatory landscape of the early developmental cell fates, we made use of three well-characterized TSC^56^, ESC^73^ and XEN cell lines^74^, that have been previously shown to be lineage-restricted, and recapitulate functional and molecular properties of their *in vivo* counterparts^56, 57, 74^ (Fig. 1a). Independent characterization of each cell line by RNA-seq analysis and immunofluorescence (IF) validated the cell-type specific expression of key signature genes, including *Cdx2, Eomes, Elf4* and *Gata3* for TSCs, *Nanog, Zfp42, Klf4* and *Pou5f1* for ESCs and *Gata4/6* and *Sox17* for XEN (Fig. 1b and Extended Data Fig.1a). PCA integrating previously published RNA-seq datasets for TSC, ESC and XEN lines (Supplementary Table 1) further confirmed that each of our samples clustered together with their respective cell type and separated from the other lineages (Extended Data Fig. 1b).

**Figure 1.**
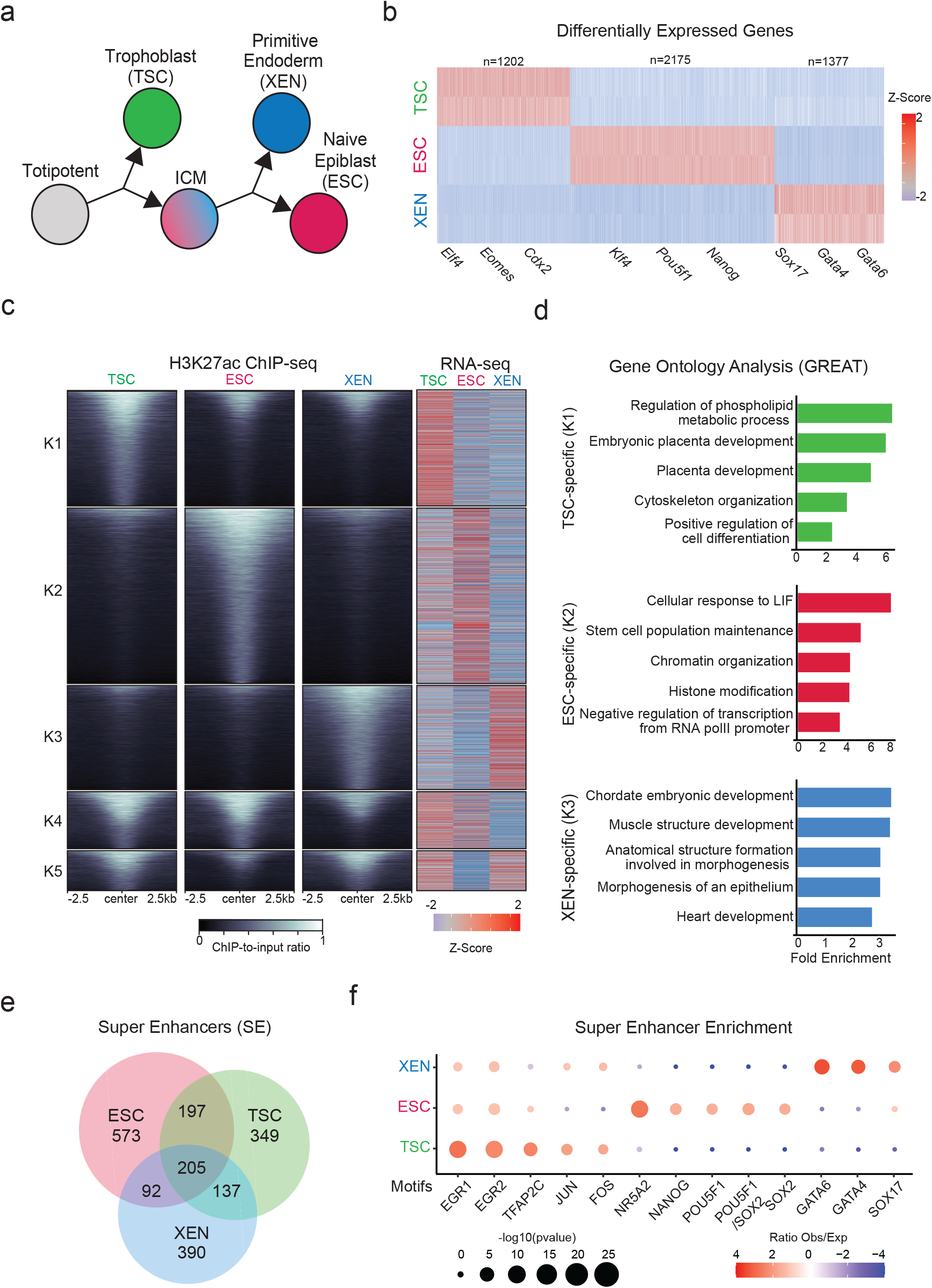
Transcriptional changes and enhancer remodeling accompany early developmental decisions. a. Schematic illustration depicting the experimental cell lines used to model early developmental fate decisions. b. Heatmap showing TSC, ESC and XEN signature genes, which are significantly upregulated in the respective cell line compared to the other two lineages (TPM>1, LogFC >2 and p-adjusted <0.01). Scale represents Z-score of normalized RNA-seq counts. RNA-seq was performed in two independent replicates for each sample. Examples of known regulators and markers of each lineage are highlighted on the bottom. For further details see also Extended Data Fig.1 c. Tornado plot (left) illustrating H3K27ac ChIP-seq signal for TSC, ESC and XEN around different clusters of peaks (+/-2.5 kb), as defined by K-means clustering (K=5) using an atlas of all H3K27ac peaks across cell lines. Scale bars denote normalized H3K27ac ChIP-seq signal over input. Heatmap (right) illustrates Z-score normalized RNA-seq levels of most proximal genes corresponding to each of the H3K27 peaks. For further details see also Supplementary Table 2. d. Gene ontology analysis (using the GREAT software) of cell type specific enhancers as identified by K-means clustering shown in (1B). Significance was calculated with the two-sided binomial test and “Region Fold Enrichment” is presented on the x-axis for selected significant (padj-value<0.05) biological processes shown in the graph. For further details see also Supplementary Table 3. e. Venn-diagram showing degree of overlap among Super Enhancers (SE) in TSC, ESC and XEN cell lines, as called by the ROSE algorithm using H3K27ac peaks as input. For further details see also Supplementary Table 2. f. Relative enrichment of TF binding motifs found in cell-type specific SE. The enrichment plots depict selected significant motifs with -log10(p-value) higher in one cell type versus the other two. Size of dots indicates the p-value (two-sided Fisher’s exact test) while color indicates the ratio of observed versus expected frequency. For further details see also Supplementary Table 3. *Note:* all statistics are provided in Supplementary Table 9.

We next performed ChIP-seq analysis for H3K27ac, which marks putative active enhancers and promoters, and ATAC-seq analysis for chromatin accessibility to map the regulatory landscapes of TSC, ESC and XEN cells. PCA clearly separated all three lineages based on either H3K27ac occupancy or chromatin accessibility (Extended Data Fig. 1b), suggesting genome-wide enhancer remodeling. K-means clustering of H3K27ac peaks across the three lineages revealed a large proportion of cell-type specific peaks (K1-K3) (Fig. 1c and Supplementary Table 2), which were predominantly located within distal intergenic and intronic regions (Extended Data Fig. 1c), while peaks shared among two or three lineages showed an overrepresentation of promoters (Extended Data Fig. 1c). As expected, the cell-type specific H3K27ac peaks were associated with elevated gene expression levels in the respective cell line (Fig. 1c). Gene ontology analysis using the GREAT tool^75^ showed that TSC-specific peaks were associated with genes involved in placenta development, XEN-specific peaks were linked to mesendoderm lineage differentiation, such as heart development, while ESC-specific peaks were associated with pluripotent stem cell maintenance and signaling, such as LIF response (Fig. 1d and Supplementary Table 3). Using the ROSE algorithm, we also identified several hundreds of Super Enhancers (SE)^76^, the majority of which were unique for each lineage (Fig. 1e Supplementary Table 2), consistent with the suggested role of SEs in cell fate regulation^69, 76–78^. Motif analysis of accessible sites within cell-type specific SE detected enrichment for known critical regulators of primitive endoderm (e.g GATA4/6 and SOX17) in XEN SE, naïve epiblast (e.g NANOG, POU5F1/SOX2, NR5A2) in ESC and trophoblast lineage (e.g TFAP2C and JUN/FOS) in TSC^67, 79–88^ (Fig. 1f and Supplementary Table 3). These results document that the distinctive transcriptional program and identity of the early developmental lineages are supported by the coordinated crosstalk of lineage-specific TFs and enhancer landscapes.

### Mapping of 3D chromatin architecture reveals multilayered genomic reorganization in early developmental lineages and complex networks of putative regulatory interactions

To investigate whether the observed remodeling of enhancer marks and chromatin accessibility among TSC, ESC, and XEN cells are also accompanied by large-scale 3D architectural rewiring, we initially performed *in situ* Hi-C (Supplementary Table 1). PCA analysis both on the level of A/B compartments (100kb resolution) and TADs (40kb resolution) clearly separated all three lineages (Extended Data Fig. 2a). Intriguingly, a higher degree of similarity was observed between TSC and XEN cells, which are both extraembryonic lineages (Extended Data Fig. 2a). Each pairwise comparison of compartment scores showed that up to 33.5% of the genome (32.5% between ESC and XEN, 33.5% between ESC and TSC and 21.1% between TSC and XEN) underwent compartmentalization changes (e.g. A-to-B, B-to-A and A or B compartment strengthening with Delta c-score >0.2 or <-0.2), albeit only ∼500-2000 genomic windows switched from A-to-B or B-to-A (Fig. 2a). In agreement with previous studies in other cellular systems,^89–91^ compartmental reorganization in TSC, ESC and XEN cells associated with transcriptional and epigenetic changes. For example, A compartment strengthening, or B-to-A switches correlated with transcriptional upregulation and gain of H3K27ac signal, while B strengthening, and A-to-B shifts associated with gene downregulation and H3K27ac loss (Fig. 2b-c and Extended Data Fig. 2b). Notably, although compartmental shifts occurred around several important developmental genes (see *Sox2* and *Foxa2* examples in Fig.2c), the majority (>80%) of cell type-specific genes and enhancers (K1/K2/K3) were not associated with compartmental changes (B-to-A). This suggests that large-scale topological changes can only explain a fraction of the extensive epigenetic and transcriptional reprogramming observed in these early developmental cell lineages. At 40kb resolution, although we observed only a few significant changes at the insulation level (<7%) between any pairwise comparison, we detected thousands (20,000-26,000) of genomic regions with significantly altered overall interactivity (within 0.5Mb window), especially when comparing ESCs with either of the extraembryonic lineages (Fig. 2d and Extended Data Fig. 2c). Gain or loss of interactivity associated with gain or loss of enhancer and transcriptional activity (Fig.2e and Extended Data Fig. 2d), respectively, documenting a rather extensive 3D chromatin reorganization that occurs along with enhancer remodeling.

**Figure 2.**
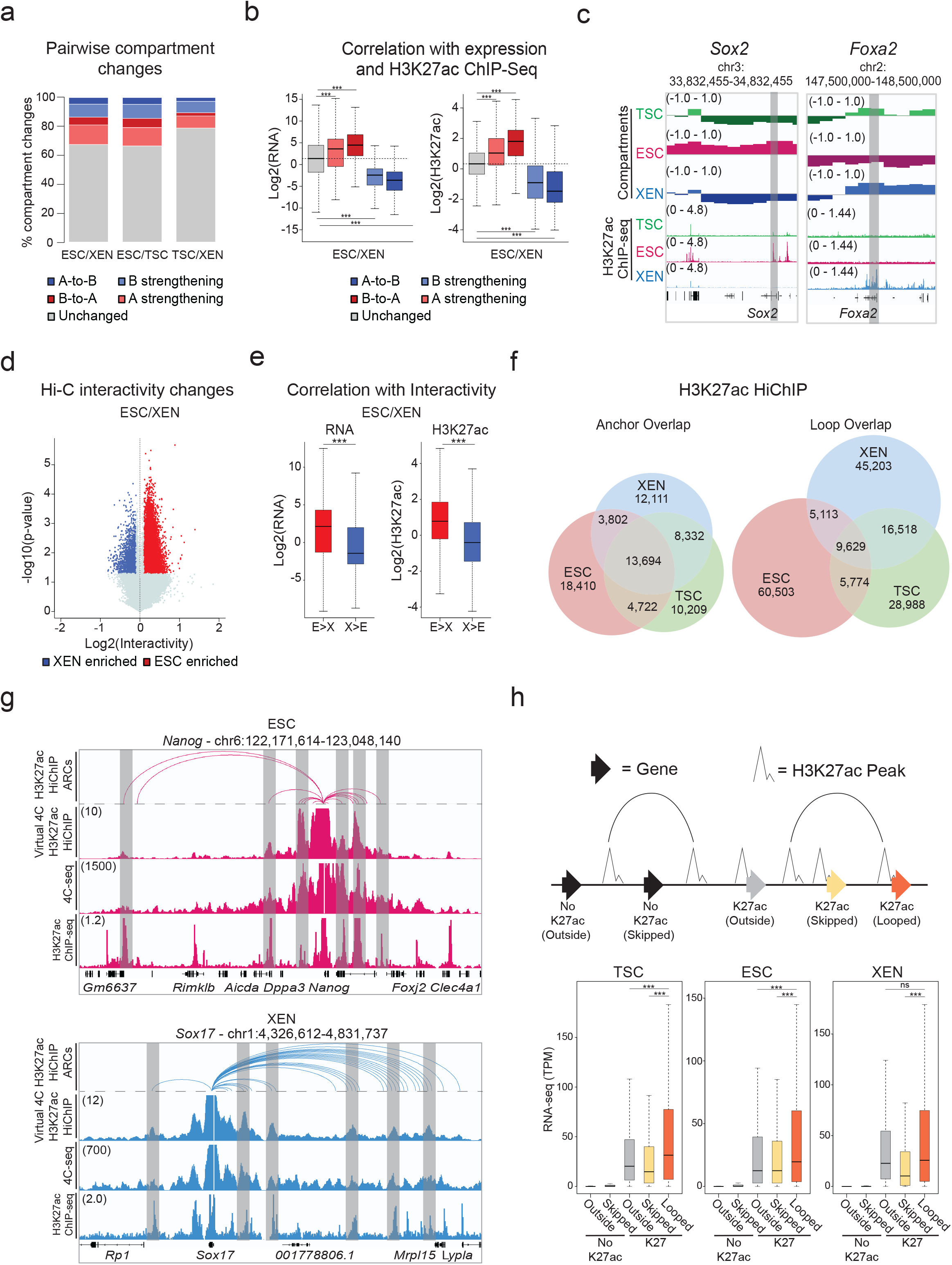
Hi-C and H3K27ac HiChIP reveals multilayered 3D genomic reorganization and complex networks of putative regulatory interactions in TSC, ESC and XEN. a. Stacked barplots showing the percentages of A/B compartment changes as detected by Hi-C for every pairwise comparison between ESC, XEN and TSC. Compartment changes at 100kb resolution were assigned to one of five groups based on their A or B status (positive or negative C-score values, respectively) in each cell type and the C-score difference between two cell lines A-to-B shifts (dark blue), B-to-A shifts (dark red), A-strengthening (light red), B-strengthening (light blue) or unchanged (grey). See Methods for details. b. Boxplots showing median expression changes (left) or H3K27ac ChIP-seq changes (right) between ESC and XEN cells at gene loci assigned to different compartment groups as described in (a). See Extended Data Fig. 2b for the other pairwise comparisons. c. Examples of A/B compartment switches around developmentally relevant genomic loci such as *Sox2* gene and SE (ESC-signature gene-left panel) and *Foxa2* (XEN-signature gene, right panel). Compartment tracks indicate c-scores, while H3K27ac tracks show normalized ChIP-seq signals. d. Volcano plot showing differential Hi-C interactivity at 40kb resolution between ESC and XEN. X-axis shows the difference of the interactivity levels, while y-axis shows -log10(p-value) as calculated by two-sided Student’s t-test. Significant changes (p-value<0.05 and Diff>0.1 or <-0.1) are highlighted in blue (gained in XEN) or red (gained in ESC). See Extended Data. Fig. 2c for the other pairwise comparison. e. Boxplots showing changes in gene expression (left) and H3K27ac ChIP-seq (right) between ESC and XEN at regions that underwent interactivity changes as described in (d). See Extended Data. Fig. 2d for the ESC and TSC pairwise comparison. f. Venn diagrams showing the numbers of shared and unique annotated anchors (left) and loops (right) in TSC (green), ESC (red) and XEN (blue) cells as detected by H3K27ac HiChIP experiments. Interactions were identified by FitHiChIP 2.0 at a 5kb resolution. g. IGV tracks showing the concordance between H3K27ac HiChIP results (presented as Arcs on top and as virtual 4C of normalized H3K27ac HiChIP signal in the middle) with independent *in situ* 4C-seq experiments around selected viewpoints (*Nanog* promoter on top and *Sox17* promoter at the bottom) along with the respective H3K27ac ChIP-seq tracks. Examples of interactions that are identified both by HiChIP and 4C-seq are highlighted in grey. The average 4C-seq signals and the H3K27ac ChIP-seq were normalized to the sequencing depth derived from two biological replicates. h. Schematic (top) defining different gene categories based on their position relative to HiChIP loops (looped, skipped, outside) and the presence or absence of promoter H3K27ac peaks (noK27ac vs K27ac). Boxplot (bottom) depicting the median gene expression levels for all gene categories in ESCs (“Outside-no K27ac” = 8110, “Skipped-noK27ac” = 5589, “Outside-K27ac” = 1129, “Skipped-K27ac = 894, “looped-K27ac” = 11020). Asterisks indicate significant differences (p-val<0.001) by Wilcoxon rank test. *Note:* all statistics are provided in Supplementary Table 9.

Encouraged by the 3D interactivity changes detected by Hi-C, we next performed H3K27ac HiChIP^43^ generating more than 2 billion reads in order to profile putative enhancer interactions in TSC, ESC and XEN cells at high genomic resolution (Supplementary Table 1). All samples passed quality control metrics validating the efficiency of HiChIP library preparation^92^ and generated more than 400 million valid pairs. By applying FitHiChIP 2.0^93, 94^ at 5kb resolution with FDR<0.05 on all datasets, we called ∼60,000-80,000 high-confidence interactions that occurred between ∼35,000-40,000 anchors in each cell type (Fig. 2f), reflecting the fact that many genomic regions engage in more than one chromatin contact. Despite the large fraction of shared anchors, we observed a poor overlap (12-16%) of chromatin interactions (“loops”) (Fig. 2f, right Venn diagram), in agreement with the high degree of regulatory rewiring indicated by Hi-C analysis. To independently validate the HiChIP called loops, we confirmed their enrichment in recently published Micro-C data in mouse ESCs^39^ by aggregate plot analysis (Extended Data Fig. 2f). Moreover, we performed high-resolution *in situ* 4C-seq analysis around enhancers and promoters of select cell-type specific genes (e.g., *Sox17* for XEN and *Nanog* for ESC), which showed high concordance both with the virtual 4C of HiChIP and the called HiChIP contacts in the respective cell type (Fig. 2g and Extended Data Fig. 2g).

HiChIP-detected interactions occurred over a large range of distances (ranging from 10kb to 2Mb) (Supplementary Table 4) with a similar size distribution among lineages (Extended Data Fig. 2e), often skipping multiple neighboring genes and enhancers, or even crossing TAD boundaries (Supplementary Table 4). Importantly, genes whose promoters engaged in at least one HiChIP contact showed significantly higher expression levels compared to not-looped genes (whose promoters were either skipped or entirely outside of loops) (Fig. 2h) in the respective cell type. Elevated expression levels of looped genes were also detected when we focused our comparison on looped and skipped genes with similar H3K27ac signal on their promoters (Extended Data Fig. 2h). This result supports the notion that H3K27ac-HiChIP contacts likely represent active regulatory interactions in all three lineages that enhance transcriptional levels of engaged genes in a targeted manner.

### 3D “hubness” associates with level, cell type-specificity and coregulation of gene expression

The positive association between looping and gene expression suggests that engagement of promoters in multiple chromatin contacts should further enhance their transcriptional output. Indeed, when we ranked promoters into quantiles based on their connectivity or “hubness” (number of distinct HiChIP-detected contacts per anchor) (Fig. 3a), we observed that higher hubness associated with progressively higher transcriptional levels (Fig. 3b) (Spearman correlation: TSC=0.35, ESC = 0.31, XEN=0.32). These observations were true across all cell lines under investigation and suggest a potential additive regulatory impact of multiple connected anchors. When we focused on the comparison of top 10% highly connected anchors (Q10) with the least connected ones (Q1) in each lineage, we found that genes with the highest promoter connectivity not only had significantly higher transcriptional levels (as shown in Fig. 3b), but also showed a strong preferential enrichment for gene ontology categories linked to either housekeeping processes or to lineage-specific functions (Fig. 3c and Supplementary Table 3). In agreement, TSC, ESC or XEN signature genes (as defined in Fig. 1b) engaged in a significantly higher number of 3D interactions in the respective cell type (Fig. 3d). We found loci encoding known master regulators among the top connected genes in each cell type, including *Klf4* in ESC (n=15 contacts) (Fig. 3e), *Gata6* in XEN (n=27 contacts) and *Cdx2* in TSC (n=26 contacts) (Extended Data Fig. 3a), suggesting that multiple regulatory contacts contribute to their robust and cell-type specific expression. Q10 anchors in ESC showed a strong and preferential enrichment for genes that were recently identified as essential for ESC survival and proliferation by two independent CRISPR screen studies^95, 96^ (Extended Data Fig. 3b). These results highlight that genes critical for survival or cell identity tend to establish multiple regulatory connections, which might act in either a cooperative or redundant fashion to ensure tight regulation and robust expression.

**Figure 3.**
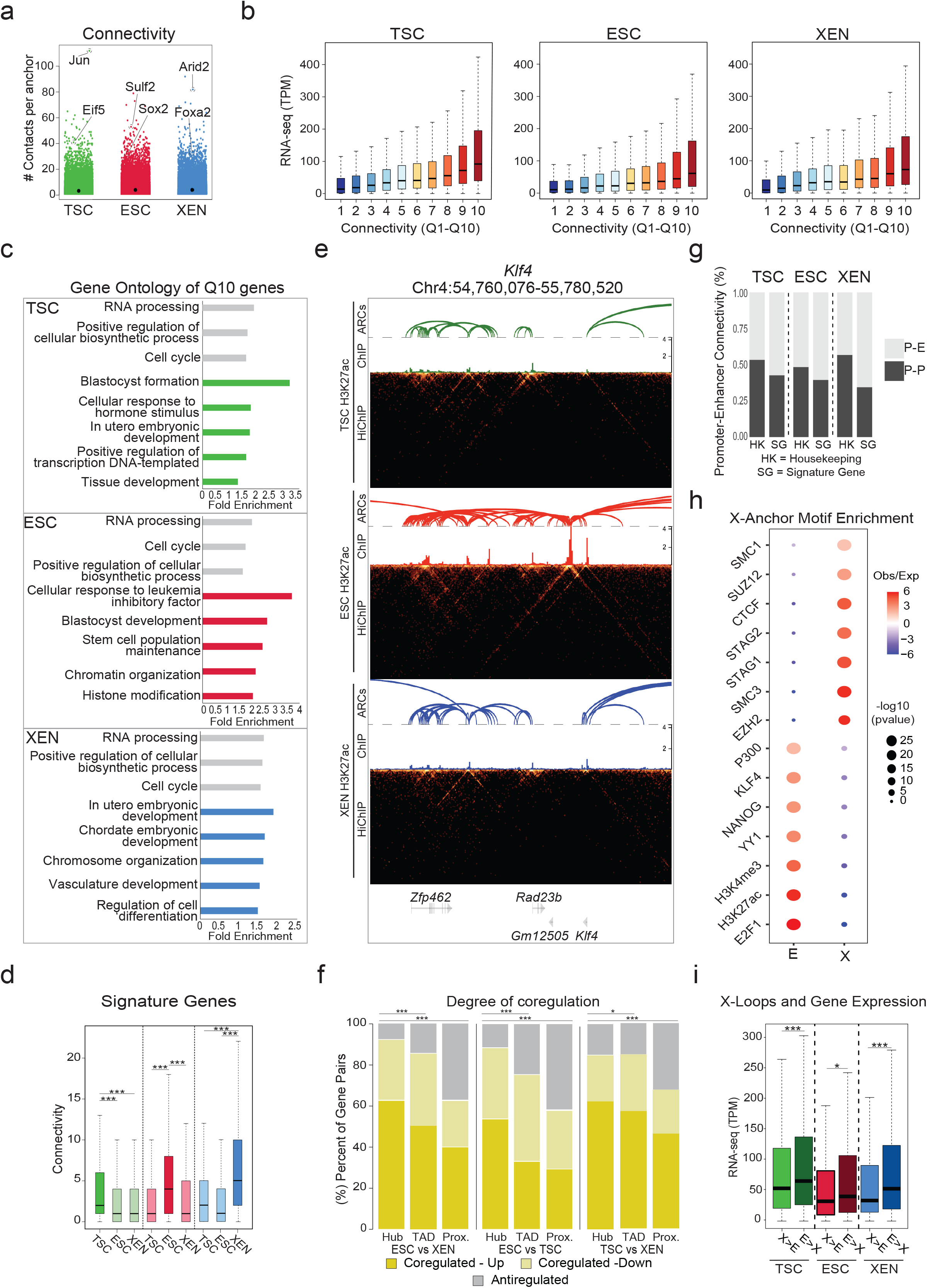
Association of high 3D hubness with levels, cell-type specificity and coregulation of gene expression in early embryonic fates. a. Plot showing the number of high-confidence HiChIP-detected contacts around each 5kb anchor (connectivity or hubness) in TSC, ESC and XEN cells. Examples of lineage-specific genes at highly connected anchors are highlighted. b. Boxplots showing median expression levels of genes with increasing HiChIP connectivity. All identified looped genes were separated into 10 quantiles based on their overall promoter connectivity, with Q1 and Q10 representing the least and the most connected. c. Gene ontology analysis showing selected housekeeping (grey) or lineage-related (colored) biological processes enriched in multi-connected Q10 genes in ESC, XEN and TSC. All genes in A compartments in the respective cell line were used as background. For further details see also Supplementary Table 3. d. Boxplots depicting the distribution and median connectivity of signature genes in each of the respective cell types. Dark colors indicate the origin of signature genes (TSC n=892 (green), ESC n=1663 (red) and/or XEN n=999 cells (blue) after removing genes with no detected loops. e. HiGlass visualization of a highly connected ESC-associated hub at the *Klf4* genomic locus shown in TSC, ESC, and XEN along with corresponding H3K27ac HiChIP-derived arcs and H3K27ac ChIP-seq signals. Interacting scores are presented in 5kb resolution. f. Stacked barplots showing the percentage of gene pairs that are coregulated (either both upregulated or both downregulated with log2 fold change>1 or <=1 and p.adj<-0.01) or anti-regulated (one upregulated and one downregulated) when comparing ESC vs XEN, TSC vs ESC and TSC vs XEN cells. Gene pairs were selected either within the same hub (connected to the same anchor by HiChIP contacts), the same TAD or in nearest linear proximity. Statistics were calculated by two-sided Fisher’s exact test (Supplementary Table 9). g. Barplots showing the percentage of Promoter-Enhancer (PE) or Promoter-Promoter (PP) pairs at housekeeping (HK) genes or signature genes (SG) in each cell type. h. Relative enrichment of TF binding motifs in either Enhancer (E) or X-linked anchors (X) in ESC. All accessible regions overlapping with an E or X were used to calculate significant enrichment (with p-value<10^-5) for different protein factors based on published ChIP-seq data using the LOLA software with all accessible regions as background. Select factors with significant enrichment either on enhancer or X-anchors are depicted. Size of dots indicates the p-value (two-sided Fisher’s exact test) while color indicates the ratio of observed versus expected. For further details see also Supplementary Table 3. i. Boxplots comparing gene expression levels of genes separated into two groups based on the relative proportion of connected X versus E anchors in the indicated cell types. The ratio of Enhancer vs X anchors is >2 in E>X hubs and <0.5 in X>E hubs. *Note:* all statistics are provided in Supplementary Table 9.

In addition to the analysis of multiconnected promoter hubs, we were also interested in identifying highly interacting enhancer hubs, meaning enhancers that form contacts with multiple genes. Such hubs could indicate coordinated regulation of two or more genes during early cell fate decisions by the same enhancer, as we and other have previously shown in other cellular contexts^42, 97–99^. To test this possibility, we focused on enhancers that interact with two or more differentially expressed genes in TSC, ESC or XEN, and examined the potential concordant (Up-Up or Down-Down) or discordant (Up-Down) regulation of all gene pairs within such hubs. Our analysis revealed a significantly higher proportion of coregulated genes within hubs, when compared to gene pairs that were most proximal to one another or pairs within matched TADs (Fig. 3f). These findings highlight that 3D hubs harbor -and potentially actively control-coregulated genes. In addition, this analysis demonstrates that integration of HiChIP interactions might be superior to any other linear or 3D features (e.g., TAD organization) to predict gene coregulation.

The positive correlation between connectivity and gene expression highlights the fact that H3K27ac HiChIP mostly detects putative active regulatory interactions. Indeed, the majority of HiChIP-detected interactions connected promoters (P: anchors contained one or more TSS) and/or putative enhancers (E: anchors with one or more H3K27ac peaks, none at a TSS) (Extended Data Fig. 3c). Intriguingly, lineage-specific genes formed predominantly interactions with enhancers than promoters (Fig. 3g), highlighting the importance of distal enhancers in cell-type specific gene regulation. On the other hand, housekeeping genes had a higher proportion of P-P interactions in all tested lineages (Fig.3g), reminiscent of recently described 3D assemblies of housekeeping genes^100^. Thus, in addition to the actual connectivity/hubness of each gene, the type of contacts could also be informative for the levels or cell-type specificity of gene expression.

In addition to the P-P, P-E and E-E contacts, about ∼25-30% of the called interactions involved one anchor with neither H3K27ac signal nor a TSS (X anchors) in each cell type. Overlap of accessible regions within X or E anchors in ESC with published ChIP-seq experiments (LOLA^101^) revealed a strong and preferential enrichment of X anchors for CTCF and Cohesin binding, as well as components of the Polycomb Repressive Complex (PRC), including EZH and SUZ12 (Fig. 3h and Supplementary Table 3). Moreover, X-anchored loops spanned significantly larger distances compared to E-E, E-P and P-P interactions (Extended Data Fig. 3d). These findings support the idea that X-anchored contacts might represent either structural or repressive loops. In support of this notion, we noticed that multi-connected genes (n>3) with a higher proportion of X vs E anchors were associated with significantly lower expression levels compared to genes with higher proportion of E connections (Fig. 3i). This held true when focusing on hubs with similar total connectivity. Finally, for conserved interactions between lineages, we noticed that switches of the anchor chromatin status from X-to-E or from E-to-X associated with upregulation or downregulation of connected genes (Extended Data Fig. 3e). These results demonstrate that not all HiChIP-detected contacts associate with positive transcriptional regulation and suggest that categorization of interactions based on the features of the involved anchors might enable a better understanding of the transcriptional fine-tuning around multi-connected gene loci.

### Association of 3D rewiring with transcriptional changes reveals classes of genes with distinct sensitivity to topological changes

Our HiChIP results document extensive fine-scale 3D reorganization during early embryonic decisions, which we independently validated for select loci by 4C-seq analysis (Fig. 4a). To determine the degree to which 3D rewiring associates with transcriptional changes, we generated an atlas of all promoter-centric contacts across the three lineages and plotted differential HiChIP connectivity vs differential RNA-seq levels between any pair of early embryonic cell types (Fig. 4b, Extended Data Fig. 4a). In every pairwise comparison, we observed a concordance of expression changes with 3D connectivity remodeling (R=0.422 for ESC/XEN, 0.318 for ESC/TSC and 0.367 for TSC/XEN), which was stronger than the correlation between transcriptional and compartmental changes (R= 0.214 for ESC/XEN, 0.098 for ESC/TSC and 0.126 for TSC/XEN). This means that gain or loss of specific HiChIP contacts at the promoter correlates with gene up- or down-regulation, respectively (3D-concordant). However, not all genes behaved the same way. In addition to a major gene group of 3D-concordant, we also identified gene loci that experienced significant changes in 3D connectivity but showed no transcriptional changes (termed “3D-insensitive”) (Fig. 4b. Extended Data Fig. 4A and Supplementary Table 5). Gene ontology analysis for the 3D-concordant gene set showed a strong enrichment for stem cell identity and developmental processes, such as pluripotency-associated signaling (ESC), tube morphogenesis (XEN) and placenta development (TSC) (Fig. 4c-d, Extended Data Fig. 4b-e and Supplementary Table 3). In contrast, 3D-insensitive genes strongly enriched for housekeeping processes, such as RNA processing, metabolism and cell cycle (Fig. 4c-d and Extended Data Fig. 4b-e). Different than 3D-concordant genes, 3D-insensitive loci showed constitutively high expression levels as well as stronger promoter H3K27ac and ATAC-seq signals across all cell types (Fig. 4e and Extended Data Fig. 4f). This analysis suggests that different types of genes have differential sensitivity or dependence on 3D connectivity changes in early embryonic lineages. Specifically, most cell type-specific genes alter their expression concordantly with 3D rewiring, while housekeeping genes maintain high expression levels that largely depend on their favorable promoter features and are likely saturated or unresponsive to connectivity changes.

**Figure 4.**
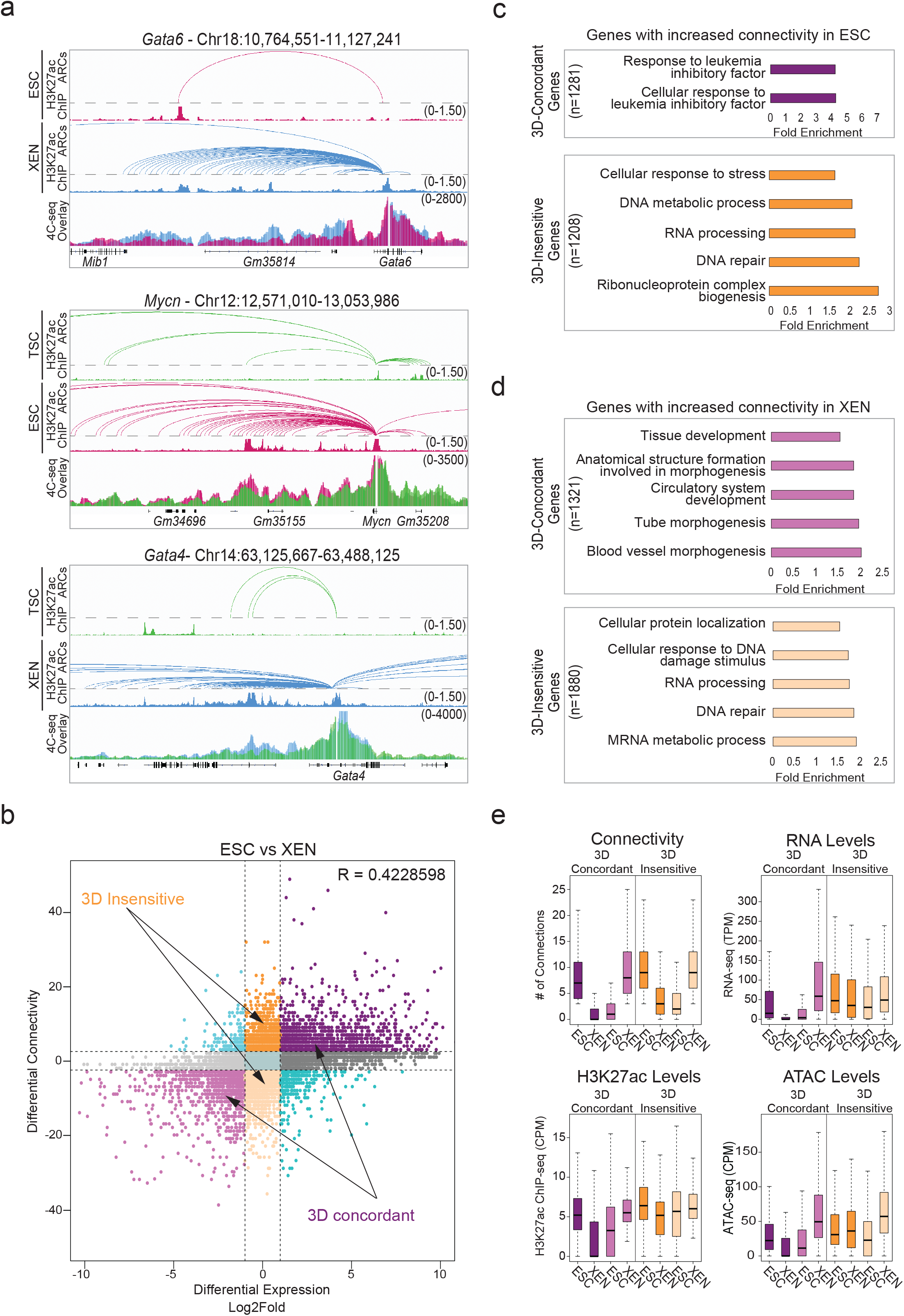
Association of 3D rewiring with cell-type specific gene expression. a. Examples of pairwise comparisons documenting 3D rewiring at developmental genes in TSC, ESC and XEN as detected by H3K27ac HiChIP (shown as arcs on top along with the respective H3K27ac ChIP-seq tracks) and validated by independent 4C-seq experiments (merged tracks at the bottom). Averaged 4C-seq signals from three biological replicates are presented after normalization to the sequencing depth. b. Correlation between differential HiChIP connectivity/hubness and differential gene expression in ESC vs XEN cells. R represents Spearman correlation identifies distinct groups of genes. We focus on the two most prominent groups: 3D-insensitive genes, defined as genes with differential connectivity >3 but no transcriptional changes (log2FC <1 or >-1) and 3D-concordant genes for which connectivity and expression changes (log2FC >1 or <-1) positively correlate (Supplementary Table 5). c. Gene ontology analysis depicting the most significant biological processes enriched in the 3D concordant (purple) and 3D insensitive (orange) groups in ESC cells as defined in (b). All genes in A compartments were used as background. For further details see also Supplementary Table 3. d. Same as in (c), but for genes with increased connectivity in XEN. For further details see also Supplementary Table 3. e. Comparison of connectivity, gene expression levels (TPM) as well as H3K27ac and ATAC CPM levels on the promoters of 3D-concordant and 3D-insensitive genes in ESC and XEN cells. Insensitive genes show higher levels of connectivity, H3K27ac, ATAC and expression in both cell types. Wilcoxon rank sum test was used for all comparisons (Supplementary Table 9). *Note:* all statistics are provided in Supplementary Table 9.

### Predictive gene expression modeling using 3D chromatin features outperforms promoter- or 1D-based models

So far, our analyses established strong links between 3D connectivity and transcriptional regulation, but also identified notable exceptions. Therefore, we sought to systematically investigate which 3D features were most important for predicting transcriptional output, including cell-type specificity and absolute expression levels. To this end, we built an optimized Random Forest machine-learning model, which we coined 3D-HiChAT, that utilizes 1D-information extracted from our ATAC-seq and H3K27ac ChIP-seq datasets and 3D-information from our HiChIP analyses (Fig.5a). Specifically, we generated a list of ten 1D, 3D or composite variables originating either from gene promoters (5kb anchor containing the TSS) or their interacting anchors-enhancers (Supplementary Table 6). After applying recursive feature selection method to eliminate features with low importance, we nominated eight predictive features (Extended Data Fig. 5a), that individually showed variable correlations with gene expression (ranging from 0.17-0.58) (Extended Data Fig. 5b). In parallel, we constructed models that only utilize 1D-information from ChIP-seq and ATAC-seq either only from the promoter region (“Promoter-centric model”) or from the extended linear neighborhood (“Linear proximity models” n=25 ranging from 10kb to 2Mb distance from promoter) for comparison with 3D-HiChAT (Fig. 5a). Random Forest classification or regression methodology was used with each of these models to predict either top 10% or bottom 10% expressing genes (classification) or absolute gene transcription levels (correlation) in each cell type, respectively. By focusing on genes with at least one HiChIP interaction in any of the three cell types, we performed Leave One Chromosome Out (LOCO) methodology to train our data in TSC for all chromosomes but mitochondrial (chrM) and chromosome Y (chrY) (n=20, chr1-19 & chrX) prior to testing on the rest of the chromosomes and cell lines.

**Figure 5.**
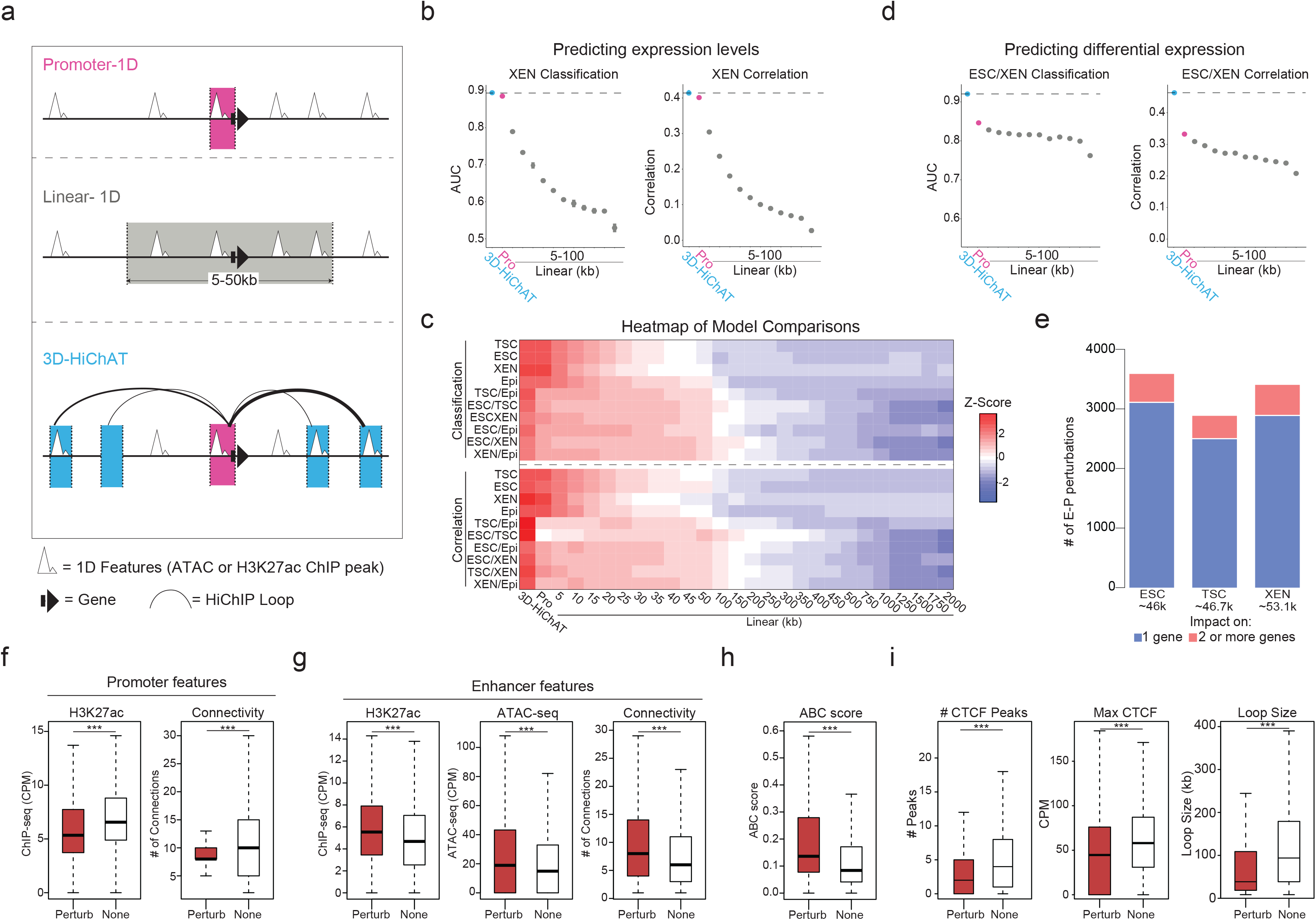
Predictive modeling using 3D chromatin features outperforms promoter- or 1D-based models for gene expression levels or cell-type specificity. a. Schematic illustration of the 1D or 3D variables used for modeling gene expression. For further details see also Supplementary Table 6. b. Area Under Curve (AUC) scores and Spearman Correlation scores generated for predicting classification of gene expression (top 10% high vs low expressing genes, left graph) and absolute levels (right graph) in XEN cells using each of our 3D-HiChAT, Promoter-1D and Linear-1D models across various distances from the TSS (5kb-100kb). Each dot represents the average score across all 20 chromosomes using the LOCO approach, while error bars show standard deviation. See also Extended Data Figure 5 for the rest of the cell lines and comparisons. For further details see also Supplementary Table 6. c. Top: Heatmap of z-scored normalized AUC values across all tested models for classification of gene expression (top 10% high or low) in each cell line or classification of differential expression (top 10% up- or downregulated) in each pairwise comparison. Bottom: Heatmap of z-scored normalized Spearman correlation values across all models for prediction of gene expression levels in each lineage or prediction of expression fold change in each pairwise comparison. Models are labeled on the bottom of the heatmap, starting from our 3D-HiChAT model, promoter and linear models ranked by distance from TSS. For further details see also Supplementary Table 6. d. Area Under Curve (AUC) scores and Spearman Correlation scores generated for predicting differential expression classification (top 10% up or downregulated, left) and fold change expression (right) between XEN and ESCs using each of our 3D-HiChAT, Promoter-1D and Linear-1D models across various distances from the TSS (5kb-100kb). Each dot represents the average score across all 20 chromosomes using the LOCO approach, while error bars show standard deviation. See also Extended Data Figure 5 for the rest of the cell lines and comparisons. For further details see also Supplementary Table 6. e. Barplots showing the numbers of E-P pairs that were predicted to reduce the expression of one (blue) or more target genes (pink) based on *in silico* perturbations in each of the cell lineages using our 3D-HiChAT model. The total number of interrogated E-P pairs for each cell type are indicated in parentheses. The distributions of predicted scores is shown in Extended Data Fig. 5f. f. Boxplots showing median H3K27ac signals (left) or Connectivity (right) at promoter anchors within E-P pairs that were predicted to be perturbed (Perturb) compared to matched number of E-P pairs that were predicted to remain unaffected (None) based on the in silico perturbations described in (e). Asterisks indicate significance pval<0.001 by Wilcoxon rank test. Although the results shown are from our ESC analysis, similar trends were detected in all cell types. (g-h). Boxplots showing median H3K27ac signal, ATAC-seq signal, Connectivity (g) and ABC score (h) at the enhancer anchors within E-P pairs that were predicted to be perturbed (Perturb) compared to matched number of E-P pairs that were predicted to remain unaffected (None) based on the *in silico* perturbations described in (e). Asterisks indicate significance pval<0.001 by Wilcoxon rank test. Although the results shown are from our ESC analysis, similar trends were detected in all cell types. i. Boxplots showing median numbers (#) and max intensities of intervening CTCF peaks as well as genomic distance (loop size) between the predicted perturbed E-P anchors compared to the non-perturbed ones, based on the *in silico* perturbations described in (e). Asterisks indicate significance pval<0.001 by Wilcoxon rank test. Although the results shown are from our ESC analysis, similar trends were detected in all cell types (Supplementary Table 9). *Note:* all statistics are provided in Supplementary Table 9.

When we tried to predict classification of gene expression (high vs low) in each cell type, we noticed that the Promoter-centric model performed very well (Area Under Curve or AUC ranging from 0.88-0.92 across all cell types), while Linear proximity models showed drastically lower accuracy when information from distal regions (>10kb) was included (Fig. 5b and Extended Data Fig. 5c). Interestingly, 3D-HiChAT consistently outperformed the promoter-centric model, albeit by a small margin (AUC up to 0.89-0.93) (Fig. 5b,5c, Extended Data Fig.5c and Supplementary Table 6). Therefore, although the epigenetic features of gene promoters are largely sufficient to explain transcriptional output, incorporating 3D features specifically from distal interacting elements rather than from the extended linear neighborhood can improve our understanding of gene expression. Notably, the same conclusions were reached when we applied Random Forest regression analysis for predicting absolute transcriptional levels (instead of classification to high or low expressing genes) (Fig. 5b and Extended Data Fig.5c) where 3D-HiChAT outperformed both promoter and linear 1D models (Spearman Correlation coefficient for Promoter-centric models 0.40-0.46 vs 3D 0.42-0.49). Importantly, 3D-HiChAT model showed similar performance and accuracy across different cell lines and species using published HiChIP, ATAC-seq and RNA-seq datasets^42^, suggesting that it is stable and generalizable (Extended Data Fig 5d).

Next, we used similar methodology (see Methods for details) to test and compare the ability of our models to predict differential gene expression among the three embryonic lineages. To avoid using the same cell lines both for training and testing, which could result in overfitting, we generated RNA-seq, ATAC-seq, H3K27ac ChIP-seq and HiChIP from a fourth embryonic cell type, mouse Epiblast Stem Cells (EpiSCs)^57^, using same methods and QC standards. The models were trained using the LOCO approach on TSC versus EpiSC data prior to testing in all other pairwise lineage comparisons using the same eight predictive features shown in Extended Data Fig. 5a. Remarkably, both classification and regression analysis demonstrated a clear superiority of the 3D-HiChAT model over promoter-centric or Linear proximity models in predicting differential gene expression (Fig.5c-d and Extended Data Fig. 5e). Promoter-based models showed poor overall predictability, highlighting that promoter information is insufficient to explain/predict cell-type specific gene expression. (Fig.5c-d and Extended Data Fig. 5e). These results highlight the importance of distal regulatory elements in cell-type specific gene expression and demonstrate that HiChIP features can enable accurate prediction of context-specific transcriptional output.

Encouraged by these results, we next used the 3D-HiChAT model to predict the relative regulatory impact of each putative enhancer on multiconnected (n>2) genes in each cell line by performing genome-wide *in silico* perturbations. Specifically, we predicted the degree of expression changes (% of perturbation) for each target gene after systematically removing each connected anchor-enhancer and recalculating all variables. E-P pairs were ranked based on their perturbation scores (%) in each cell line separately and cut-offs (for high-confidence perturbation) were determined at the points where the slope of the tangent along the curve exceeded the value of one (Extended Data Fig. 5f). Although we observed perturbations in both directions (positive and negative perturbation), we focused specifically on perturbations that caused gene downregulation, suggesting a putative enhancer function. Using this strategy, we identified ∼4,300 out of the 46,000 interrogated E-P pairs that passed the cut-off (<-9.91%) in ESCs, ∼3,400 out of 46,700 E-P pairs in TSC (< −12.55%) and ∼4,200 out of 53,100 in XEN (< −11.20%) (Fig. 5e and Extended Data Fig. 5f).

To gain more insights into the features that determine the degree of susceptibility or resistance to expression changes upon *in silico* perturbation, we directly compared the predicted functional enhancer-promoter pairs (Perturb) with an equal number of non-perturbed ones (None). Genes within the perturbed group were characterized by significantly lower ChIP-seq signal at their promoters as well as lower overall promoter connectivity compared to non-affected genes (Fig. 5f), suggesting that high promoter activity, and/or a high number of contacts could compensate for the loss of a single anchor. This aligns with our analysis about the 3D-insensitive gene set that appear irresponsive to connectivity changes (Fig. 4e). On the other hand, anchors predicted to perturb gene expression-compared to the non-perturbing ones-had significantly stronger H3K27ac signal and contact probabilities (Fig. 5g), in agreement with the recently published Activity-By-Contact (ABC) model^46^. Interestingly, although the 3D-HiChAT predictions showed a good correlation with ABC scores, (R=-0.40795) (Extended Data Fig. 5g) with most of the high-ABC enhancers showing also high 3D-HiChAT perturbation scores (Fig. 5h), we also observed several enhancers with high 3D-HiChAT scores but low ABC. These enhancers were at higher distances (median = 50kb / mean=90.75 kb) compared to the ones with high ABC (median = 15kb / mean = 20.47 kb), suggesting that our model might be able to capture more distal functional enhancers (Extended Data Fig. 5h). Nevertheless, comparison between the Perturb or None groups according to 3D-HiChAT showed that predicted impactful enhancers were significantly closer to their target genes and crossed significantly fewer and weaker CTCF binding sites (Fig. 5i). This is consistent with the notion that functional enhancers reside within the same insulated neighborhood or TAD with their target genes^30, 34, 102, 103^ although we predicted a small fraction (589/42331=13.92%) of impactful enhancers that crossed TAD boundaries.

Finally, we made an intriguing observation that the predicted impactful enhancers were also characterized by significantly higher hubness (Fig.5g), supporting the notion that enhancer 3D connectivity could indicate stronger regulatory impact and reflect a more central position in regulatory networks. This finding might also suggest that multiconnected enhancers might have regulatory impact on more than one gene, operating as 3D regulatory hubs. In total, 3D-HiChAT identified 484 enhancer hubs in ESC (controlling 1108 genes), 392 hubs in TSC (controlling 904 genes) and 523 hubs in XEN (controlling 1317 genes) whose deletion predicted downregulation of at least two up to eight different genes (Supplementary Table 6) (Fig.5e).

### Experimental validations of the 3D-HiChAT model reveal novel functional enhancers and hubs in ESC and XEN cells

The above-mentioned results suggest that 3D genomics data generated in TSC, ESC and XEN cells, combined with the 3D-HiChAT model, could enable discovery of new core enhancers that dictate these early cell fates. To experimentally test this, we initially focused on a complex, hyperconnected locus in ESCs that spans ∼1.3Mb and harbors, among others, two important genes implicated in maintenance or acquisition of pluripotency *Tfcp2l1* and *Gli2*^104–109^. According to our HiChIP results, both genes reside in the same A compartment in ESCs and form connections with a total of 17 proximal and distal putative enhancers, which show variable perturbation scores based on 3D-HiChAT (Fig. 6a-b and Extended Data Fig. 6a). Among them, we decided to experimentally test two shared putative enhancers, *Enh3* and *Enh14*, of which *Enh3* is predicted to only control *Tfcp2l1* while *Enh14* has predicted regulatory impact on both genes. To experimentally test these predictions, we transduced an ESC line stably expressing dCas9-BFP-KRAB (CRISPRi) with guide RNAs that target each of the shared enhancers or the gene promoters (Extended Data Fig. 6b). After transduction and selection (n≥3 independent experiments per gRNA), RT-qPCR was used to determine impact on gene expression compared to an empty vector control. In agreement with our predictions, CRISPRi silencing of *Enh3* caused significant downregulation of *Tfcp2l1* only (Extended Data Fig. 6b), while silencing of *Enh14* significantly reduced the expression of both *Tfcp2l1* and *Gli2* (Fig.6c-e). The concordant downregulation of both enhancer-connected genes supports its function as a 3D regulatory hub. Intriguingly, CRISPRi-mediated silencing of *Enh14* had no significant impact on other connected genes, in agreement with the lower 3D-HiChAT predicted perturbation scores on these genes.

**Figure 6.**
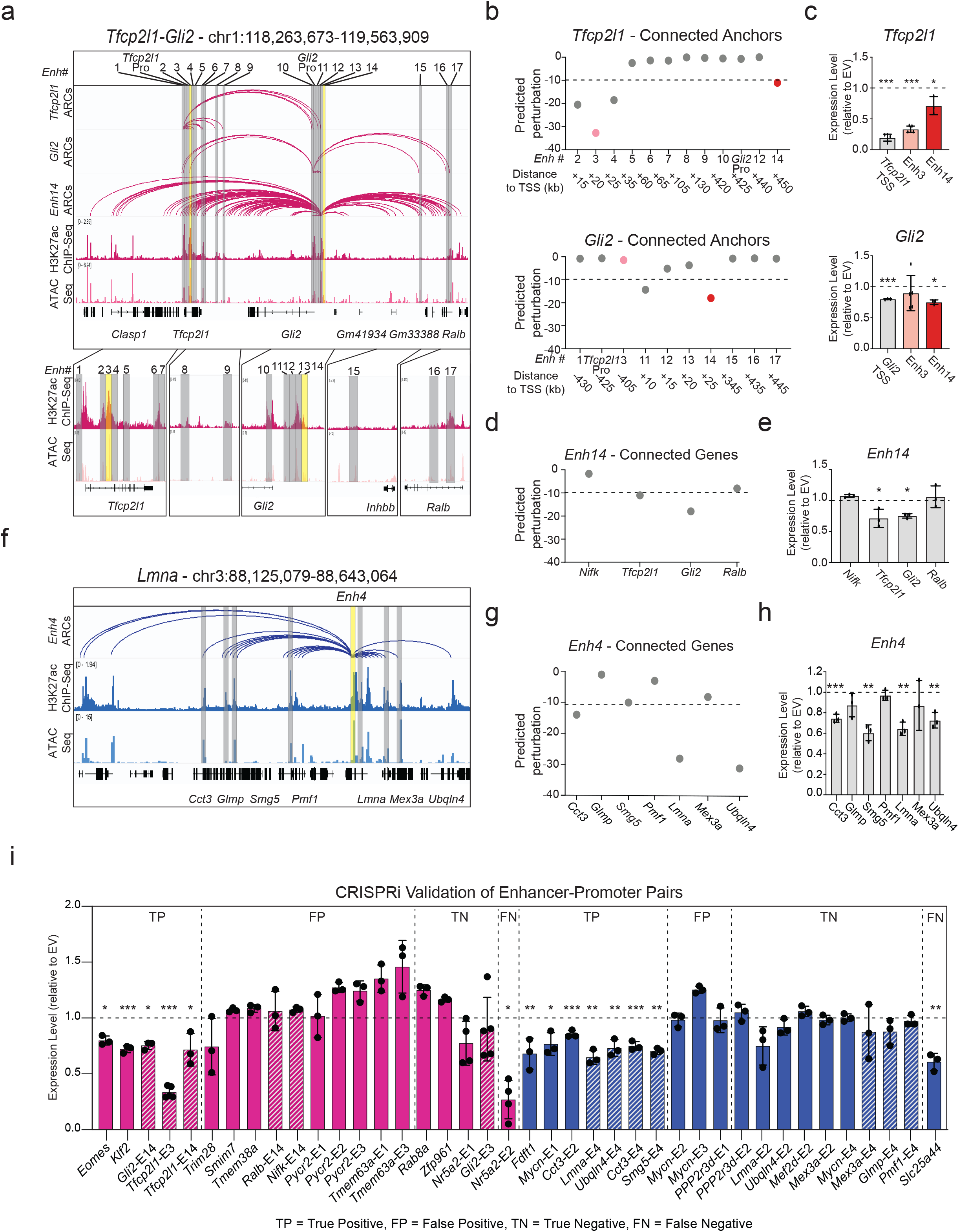
Experimental validation of predicted enhancers in ESC and XEN. a. IGV tracks of the *Tfcp2l1-Gli2* locus depicting putative regulatory elements that contact either one or both genes in ESCs. H3K27ac HiChIP derived arcs originating from each gene promoter or *Enh14* are shown along with H3K27ac ChIP-Seq and ATAC-seq peaks. Grey bars highlight all putative enhancers, while yellow bars indicate enhancers targeted by CRISPRi in this study. b. Predicted perturbation scores generated by 3D-HiChAT for each putative enhancer connected to the *Tfcp2l1* or *Gli2* promoters. The dotted line indicates the cut-off (<-9.9) we chose for potentially impactful hits in ESC (see Extended Data Fig.5f and Methods). Colored dots refer to enhancers targeted by CRISPRi. c. Relative mRNA levels of *Tfcp2l1 or Gli2* upon CRISPRi-targeting of the indicated regions compared to control cells infected with empty vector (EV). Dots indicate biological replicates (n≥3 independent experiments). Error bars indicate mean ± SD. Statistical analysis was performed by one-tailed unpaired student t-test. Asterisks indicate significance < 0.05. d. 3D-HiChAT-based predicted perturbation scores for genes connected to *Enh14*. The dotted line indicates the cut-off (<-9.9) we chose for potentially impactful hits in ESC (see Extended Data Fig.5f and Methods). e. Relative mRNA levels of Enh14-connected genes upon CRISPRi perturbation of Enh14 in ESCs compared to control cells infected with empty vector (EV). Dots indicate biological replicates (n=independent experiments). Error bars indicate mean ± SD. Statistical analysis was performed by one-tailed unpaired student t-test. Asterisks indicate significance < 0.05. f. IGV tracks of the depicting a highly connected enhancer hub (Enh4 shown in yellow) which interacts with 7 gene promoters (shown in grey) in XEN. HiChIP arcs originating from Enh4 as well as H3K27ac ChIP-Seq and ATAC-seq are shown. g. Predicted perturbation scores for genes connected to *Enh4* hub. The dotted line indicates the cut-off (<-11.20) we chose for potentially impactful hits in XEN (see Extended Data Fig.5f and Methods). h. Relative mRNA levels of *Enh14*-connected genes upon CRISPRi targeting of *Enh14* in XEN compared to control cells infected with empty vector (EV). Dots indicate biological replicates (n= 3independent experiments). Error bars indicate mean ± SD. Statistical analysis was performed by one-tailed unpaired student t-test. Asterisks indicate significance < 0.05. Of note, *Smg5* scored borderline below the cut-off (−10.61), but was still validated experimentally. i. Barplots summarizing the expression changes upon CRISPRi experimental perturbations of a total of 40 enhancer-promoter pairs in ESC (Pink) and XEN (blue), that were either predicted to be positive hits (cut-off <-9 for ESC and <-11.2 in XEN) or negative, based on the 3D-HiChAT model. Each bar shows the mean RT-qPCR values for each gene upon CRISPRi targeting of their candidate enhancers relative to the values in the Empty Vector (EV) control cells and after normalization relative to housekeeping genes (*Hprt* for ESC and *Gapdh* for XEN). For some genes, multiple enhancers were tested as indicated in the title (see Supplemental Table 7 for details). Shaded bars indicate that these data are also presented in the context of their respective hubs in the (c), (e) or (h) panels. Dots indicate biological replicates (n=3 independent experiments). Error bars represent mean ± SD. Statistical analysis was performed by one-tailed unpaired student t-test. Asterisks indicate significance < 0.05. Results are grouped into four categories (TP, FP, TN, FN) reflecting either the concordance or discordance between predictions and experimental validations. Of note, *Eomes* and *Smg5* scored borderline below the cut-off (−9.8 and −10.61), but was still validated experimentally *Note:* all statistics are provided in Supplementary Table 9.

By establishing a similar CRISPRi system in XEN cells (Extended Data Fig. 6c) we were able to also validate a novel enhancer hub (*Enh4*) connected to 7 genes (including *Cct3*, *Glmp*, *Smg5*, *Pmf1*, *Lmna*, *Mex3a* and *Ubqln4*) across a 520kb region (Fig.6f) with different predicted impact on each gene (Fig. 6g). CRISPRi-mediated targeting of this hub led to significantly downregulated levels of *Lmna*, *Cct3, Smg5* and *Ubqln4,* while other connected genes (*Glmp*, *Pmf1* and *Mex3a*) remained unaffected (Fig. 6h), in agreement with our model predictions.

Encouraged by these results, we extended our experimental perturbations to a total of 40 enhancer-promoter pairs in ESC (n=20, pink) or XEN (n=20, blue), which were selected to represent loci with moderate connectivity (between 2-12 connections) and variable 3D-HiChAT perturbation scores (ranging from −0.02 to −46.8) (Fig. 6i and Supplementary Table 6). Our experiments revealed 12 true positive hits (including novel enhancers around important developmental genes such as *Klf2, Eomes* and *Mycn*) and 13 true negative hits. Ranking E-P pairs based on the perturbation scores and classifying genes as perturbed or not based on CRISPRi results showed that our model had an overall accuracy of 0.71 (Extended Data Fig. 6d). Although this is potentially an underestimation, due to the variable efficiencies of the gRNAs, it indicates that additional improvements and metrics are needed for more accurate predictions. Interestingly, more than half of our validated enhancers had very low ABC scores (<0.2), (Extended Data Fig. 6e) partly reflecting their higher distance to their target genes, suggesting that our model might be more suitable in predicting distal functional enhancers.

Together, these results demonstrate the ability of 3D-HiChAT to predict complex regulatory relationships, including enhancer hierarchies around multiconnected genes as well as enhancer-promoter specificity of multiconnected enhancers. Given the stable performance of the model across different cell types and species (see Extended Data Fig.5d), 3D-HiChAT could be applied in different biological systems to nominate candidate functional enhancers or help interpretation of disease-associated structural variants.

## DISCUSSION

Cell-type specific transcriptional programs are controlled by the activity of transcription factors and their target enhancers^110–113^. Therefore, studying the mechanisms of enhancer activity and specificity is essential for understanding and modulating the mechanisms that dictate cell fate decisions. In this study, we applied H3K27ac HiChIP and other genomics technologies to map, at high-resolution, the landscapes and 3D interactomes of putative active enhancers in the context of the first embryonic lineages and establish associations with transcriptional behavior and cell identity. Our results generated detailed 3D networks of enhancer-promoter connections in mouse TSCs, ESCs and XEN cells and provided a resource of predicted functional enhancers for each lineage as well as proof-of-concept validations. Moreover, our integrative analysis and gene expression predictive model revealed new - and potentially universal-insights into the functional interplay between 3D connectivity and transcription.

Physical proximity -but not necessarily physical contact-is considered the most likely mechanism for functional communication between genes and distal regulatory elements^102^ and an important feature for assigning enhancers to their cognate target genes^114^. In agreement with previous studies in various cellular contexts^42, 99, 115, 116^, our study revealed a strong positive correlation between 3D connectivity -or “hubness”- and gene expression across lineages, but also important exceptions which reflect the intricate nature of transcriptional regulation in the context of complex and dynamic 3D networks. Specifically, our integrative analysis and predictive modeling uncovered distinct principles and 1D/3D features that influence (i) the relative susceptibility of multi-connected genes to topological changes or enhancer perturbations and (ii) the relative regulatory impact of individual enhancers on one or more target genes. For example, we observed a strong concordance between transcriptional and topological changes around lineage-specific genes, suggesting that the *de novo* establishment (or strengthening) of long-range interactions with distal enhancers is critical for robust and context-specific activation of these genes. On the contrary, housekeeping genes appeared insensitive to 3D rewiring, suggesting that their high expression levels are likely driven from their promoters, which are saturated or irresponsive to additional regulatory input. This result aligns with recent high-throughput reporter assays that interrogated enhancer-promoter compatibility and found a reduced responsiveness of housekeeping promoters to distal enhancers^117^. Moreover, our *in silico* and experimental perturbations showed that highly connected genes -both housekeeping and developmental-tend to be less susceptible to individual enhancer deletions, suggesting functional redundancy among enhancers and phenotypic robustness in line with previous studies in different cellular contexts^118, 119^

Several computational models have been developed to predict putative functional enhancers in various cellular contexts either based on 1D features (e.g. chromatin accessibility, histone marks, TF/co-factor binding, nascent transcription etc.)^120–125^ and/or 3D features, such as CTCF binding, insulation^33, 126^ or contact probability with target genes^46, 127, 128^. These predictions become particularly challenging in the context of highly interacting hubs^129^ where multiple genes and putative regulatory elements come in spatial proximity (albeit not necessarily all at the same time and allele) making it hard to dissect which of these interactions have positive, negative or neutral regulatory impact. 3D-HiChAT predictions and functional validations show that consideration of both 1D and 3D features extracted from 3D enhancer-promoter networks enables better predictions of (i) transcriptional behaviors, such as levels and cell-type specificity of gene expression or probability of gene co-regulation and (ii) of complex regulatory relationships, including enhancer hierarchies or redundancies and enhancer-promoter specificities. Indeed, based on our predictions, we were able to identify and validate several “dominant” enhancers around multiconnected developmental genes, as well as novel functional enhancer hubs,

responsible for the coordinated regulation of more than two genes in ESC or XEN. Importantly, not all connected genes respond to the same enhancer and not all putative enhancers contributed to the regulation of their interacting genes. In agreement with previous studies, 3D-HiChAT showed that the relative contact frequency between enhancers and promoters and their putative activity/accessibility (as indicated by H3K27ac ChIP-seq and ATAC-seq) are important predictors of their regulatory relationships. However, our model also took into consideration the secondary interactions of each enhancer and showed that high degree of enhancer hubness is predictive of stronger regulatory impact upon perturbation, and potentially on multiple connected/coregulated genes. These findings nominate 3D hubness as an important predictive feature of regulatory centrality and suggest that mapping of 3D hubs could help dissect regulatory hierarchies and predict core modules (both critical genes and enhancers) that instruct cell-type-specific transcriptional programs.

Collectively, our studies showed that 3D-HiChAT is a stable model, is generalizable to different cell-types and species, performs better than 1D-based models and enables prediction of complex regulatory relationships around multiconnected genes and enhancers. However, our results also highlighted the need for further improvements in the modeling and the experimental strategy. Generation and utilization of ultra-resolution (sub-kb) 3D genomics datasets and consideration of additional variables, such as binding of CTCF or lineage-specific transcription factors or enhancer-associated co-factors, could further improve model performance. On the other hand, systematic high-throughput functional screens of putative positive and negative regulatory elements (e.g. X anchors) during dynamic cell fate transitions, will enable a deeper understanding of the regulatory relationships (hierarchies, redundancies, synergies or competitions) and inform development of better modeling approaches for prediction of core regulatory enhancers and hubs.

In conclusion, our study systematically mapped the dynamic 3D enhancer chromatin networks within the first embryonic (EPI) and extraembryonic (TE and PrE) cell fates and nominated candidate core enhancers for future high-throughput functional perturbations *in vitro* or *in vivo*. Moreover, our integrative analysis and 3D-HiChAT predictive model revealed conserved principles of transcriptional regulation through long-range interactions, providing a framework for understanding and modulating lineage-specific transcriptional behaviors.

## METHODS

### Cell culture

The feeder-dependent murine ESC line v6.5 and feeder free Bruce-4 cells were cultured in 2% gelatin-coated (SIGMA, G1393) ventilated-cap flasks, using standard serum/LIF/2i conditions in DMEM (GIBCO, 41966) supplemented with 15% fetal bovine serum (GIBCO, 10270), 1 mM sodium pyruvate (Gibco, 11360070), 2mM L-Glutamine (GIBCO, 15030), 0.1 mM non-essential amino acids (Gibco, 11140050), 100 U/ml Penicillin/100µg/ml Streptomycin (Gibco,15140163), 100 μM β-mercaptoethanol (SIGMA, 63689), 1000 U/ml leukemia inhibitory factor (derived in house), (1 μM MEK inhibitor (Stemgent, 04-0006) and 3 μM GSK3 inhibitor (Stemgent, 04-0004)^73^. TSC feeder-dependent cells were cultured on mitomycin-treated MEFs at a 40-60% density in RPMI1640 (VWR, 10-040-CV) supplemented with 20% fetal bovine serum (GIBCO, 10270), 1mM sodium pyruvate (Gibco, 11360070), 100 U/ml Penicillin/100µg/ml Streptomycin (Gibco, 15140163), 100 μM β-mercaptoethanol (SIGMA, 63689), 25 ng/ml bFGF (Thermo, PHG0360) and 1 μg/ml heparin^56^. Established XEN cells were cultured in standard XEN cell culture conditions^74, 130^. Cells were plated onto tissue culture grade plates coated with 0.2% gelatin (Millipore Sigma, G9391) in DMEM supplemented with 15% fetal bovine serum (GIBCO, 10270), 1 mM sodium pyruvate (Gibco, 11360070), 2mM L-Glutamine (GIBCO,15030), 0.1 mM non-essential amino acids (NEAA; Gibco, 11140050), 100 U/ml Penicillin/100µg/ml Streptomycin (Gibco, 15140163), 100 μM β-mercaptoethanol (SIGMA,63689). ESC and XEN cell were passaged every 2-3 days (∼70-80% confluence), while TSC were passaged every 4-5 days by washing with phosphate buffered saline (1xPBS) followed by brief incubation in 0.05% Trypsin-EDTA (Gibco, 25300054) at 37°C ∼2-3 mins). Trypsin activity was neutralized with serum-containing media (3x volume of Trypsin used) and dissociated cells were centrifuged at 300*g* for 5 mins before resuspending in culture media. Cells were replated at 1:8-1:10 dilution. Embryo derived EpiSC cells were cultured in fibronectin coated plates in DMEM-F12 (Fisher, 10-565-018), supplemented with 100 U/ml Penicillin/100µg/ml Streptomycin (Gibco, 15140163), 2 mM L-glutamine (GIBCO, 15030), 1mM non-essential amino acids (Gibco, 11140050), 50μg/ml bovine serum albumin (Gibco, 15260-037), 0.11 mM β-mercaptoethanol (SIGMA, 63689), 20ng/ml Activin A (Peprotech 120-14E), Fgf2 (12.5 ng/ml, Thermo, PHG0360) and 0.5% N2 (Thermo, 17502048) and 1% B27 supplement (Thermo, 12587010)^57^.

KH2 ESC cells were converted into EpiSC cells as previously shown ^131^. Briefly, ESCs were plated on fibronectin coated plates in 50% DMEM-F12 (Thermo Fisher Scientific, 11320033), 50% Neurobasal (Thermofisher, 21103049), 0.5% N2 (Thermo, 17502048) and 1% B27 supplement (Thermo, 12587010), 2 mM glutamax (GIBCO, 15030), 100 U/ml Penicillin/100µg/ml Streptomycin (Gibco, 15140163), and 0.1% β-mercaptoethanol (SIGMA, 63689), supplemented with 12.5 ng/ml bFGF (12.5 ng/ml, Thermo, PHG0360), 20 ng/ml Activin A (Peprotech 120-14E), and 1% Knockout Serum Replacement (Thermofisher, <u>10828010)</u>. Upon 48h EpiLCs were dissociated into small clumps (∼3-5 cells) with TrypLE (Fisher,12605010) and plated on mouse fibroblast feeders in 50% DMEM-F12, 50% Neurobasal, 0.5% N2 (Thermo, 17502048) and 1% B27 supplement (Thermo, 12587010), 2 mM glutamax (GIBCO, 15030), 100 U/ml Penicillin/100µg/ml Streptomycin (Gibco, 15140163), and 0.1% β-mercaptoethanol (SIGMA, 63689), supplemented with 12.5 ng/ml bFGF (12.5 ng/ml, Thermo, PHG0360), 20 ng/ml Activin A (Peprotech 120-14E), Wnt inhibitor (Selleck, S7238) and cultured for 2-3 days.

### Lentiviral production and infection

293T cells were transfected with overexpression constructs along with the packaging vectors VSV-g, Tat, Rev and Gag-pol using PEI reagent (PEI MAX, Polyscience, 24765-2). The supernatant was collected after 48 and 72 h, and the virus was concentrated using polyethylglycol (Sigma, P4338). Cells were infected in medium containing 5 μg ml^−1^ polybrene (Millipore, TR-1003-G), followed by centrifugation at 1300g for 90 min at 32°C.

### CRISPRi

XEN cells were infected with lentiviruses harboring the pHR–SFFV–dCas9–BFP–KRAB vector (Addgene, cat. no. 46911), while ESC v6,5 cells were infected with a modified version of the plasmid in which the SFFV promoter was replaced with an Ef1a promoter ^42^. Cells expressing BFP were selected by 3 consecutive rounds of FACS sorting (enriching only for the high expressing cells each time). The resulting, ESC stably expressing the dCas9–BFP-KRAB cells, were then infected with a lentivirus harboring the pLKO5.GRNA.EFS.PAC vector (Addgene, cat. no. 57825) containing either a single or 2 gRNAs targeting the region of interest. Due to the Purmocyin resistance the XEN-dCas9-BFP-KRAB cells were infected with a modified version of the pLKO5.GRNA.EFS.PAC vector (Addgene, cat. no. 57825) replacing puromycin with blasticidin resistance. Cells were selected with puromycin (LifeTech, K210015) or blasiticidin for 4 days and subsequently collected for RT–qPCR analysis. The guide RNAs targeting each enhancer together with the RT–qPCR primers used are described in Supplementary Table 7.

### Immunofluorescence

IF experiments were performed as previously described with a few modifications ^132^. Cells were plated on sterile glass coverslips and cultured for 24h-48h until they reached a 70%-80% confluency. Cells were fixed in freshly prepared 2% PFA/1xPBS for 10 minutes at RT, permeabilized with 0.5% v/v Triton X-100/1xPBS for 10 minutes and rinsed with 1xPBS. Cells were blocked in 1% w/v BSA/1xPBS for 30 minutes at RT, incubated with the primary antibody for one hour at RT in a dark and humidified chamber, rinsed 3 times in 1xPBS, cells were then incubated with the secondary antibody for 45 minutes at RT in a dark and humidified chamber, rinsed 3 times with 1xPBS and finally left to air-dry off water residuals. Finally, the coverslips were mounted with ProLong Gold antifade reagent supplemented with DAPI for nuclear DNA staining. IF signals were examined on a Nikon Eclipse Ti V5.20microscope unit with an Andor Zyla VSC-01979 camera, using a 20x objective and images were analyzed using Fiji Is Just ImageJ (FIJI)^133^. The following primary antibodies and their dilutions used in this study were: rabbit anti-GATA6 (Bethyl, 1:200), mouse anti-Gata-4 (Santa Cruz, 1:100), rabbit anti-NANOG (Bethyl, 1:300), mouse anti-Oct4 (Santa Cruz, 1:100), rabbit anti-Eomes (Abcam, 1:400), mouse anti-Gata3 (Santa Cruz, 1:100). Secondary Alexa Fluor-conjugated antibodies (Invitrogen) were used at a dilution of 1:500.

### cDNA synthesis and RT-PCR

For quantitative expression analysis, whole cell RNA extract was prepared using the RNeasy Mini kit (Qiagen, 741106) following the manufacturer’s instructions. In order to eliminate DNA contamination, RNA samples were treated with DNase I (Qiagen, 79256). cDNA synthesis was performed using 1ug total RNA. In parallel with reverse transcriptase reactions, control reactions devoid of the enzyme were prepared in order to verify the absence of DNA contamination in the subsequent quantitative PCR (qPCR) reactions. 2.5% of the cDNA produced was used for each qPCR reaction using the SYBR Green PCR Master mix (Life technologies, A2577) according to the manufacturer’s instructions. Real-time qPCR results were analyzed with the standard ΔΔ cycle threshold method and re sults were initially normalized to the expression of either HPRT (ESC) or GAPDH (XEN cells) followed by a second normalization to the corresponding Empty Vector that was used in each biological replicate. Statistical analysis was performed by one-tailed unpaired student t-test. Significance is indicated as: *P < 0.05, **P < 0.01 and ***P < 0.001. The primer sets used for mRNA quantitation are provided in Supplementary Table 7.

### RNA sequencing & library preparation

cDNA library for RNA sequencing (RNA-seq) was generated from 100 to 400 ng total RNA using TruSeq RNA Sample Preparation Kit (20020594) according to the manufacturer’s protocol. For each cell line 2 biological replicates were sequenced and analyzed. Briefly, poly(A)–tailed RNA molecules were pulled down with poly(T) oligo–attached magnetic beads. Following purification, mRNA was fragmented with divalent cations at 85C and then cDNA was generated by random primers and SuperScript II enzyme (Life Technologies). Second-strand synthesis was performed followed by end repair, single ‘A’ base addition, and ligation of barcode-indexed adaptors to the DNA fragments. Adapter specific PCRs were performed to generate sequencing libraries. Libraries were size-selected with E-Gel EX 2% agarose gels (Life Technologies) and purified by QIAquick Gel Extraction Kit (QIAGEN). Libraries were sequenced on an Illumina HiSeq 4000 platform on SE50 mode at the Weill Cornell Medicine Genomics Core Facility.

### ChIP-exo

ChIP-exo was performed as previously described with mild modifications^134^. Briefly, 10 million cells were used per replicate for TSC, ESC and XEN. Initially, cells were crosslinked in 1% formaldehyde at RT for 10 minutes and quenched with 125mM glycine for 5 mins at RT. Cell pellets were washed twice in 1xPBS After the final wash and centrifuge, the pellet was snap frozen before extraction. Frozen cell pellets were processed as described previously in ChIP-exo 5.0 protocol^134^. A total of 10 M cells were used per each replicate of library and 3 µg of anti-CTCF antibody (Sigma-Aldrich, 07-729) was used for o/n chromatin immunoprecipitation at 4°C. Libraries were sequenced on an Illumina NextSeq 550 platform on SR100 mode at the Cornell Ithaca Epigenomics Core Facility. ChIP-seq data have been deposited in the Short Read Archive (SRA) under the accession codes GSE212992. For further details see also Supplementary Table 8.

### H3K27ac ChIP-seq

ChIP-seq was performed as previously described^42^, with a few modifications. 10 million cells were used per replicate for TSC, ESC and XEN and *in vitro* derived EpiSC cells. Initially cells were crosslinked in 1% formaldehyde at RT for 10 minutes and quenched with 125mM glycine for 5 mins at RT. As a normalization control ^135^, 5 million formaldehyde-fixed *Drosophila* nuclei were added to each sample. Cell pellets were washed twice in 1xPBS and resuspended in 300ul lysis buffer (10mM Tris pH8, 1mM EDTA, 0.5% SDS) for at least 15 minutes. Next, chromatin was sonicated in a Pico bioruptor device for 10 cycles with the length of the intervals being 30sec on/off, in order to produce 300-800 bp chromatin fragments. Sonicated chromatin was then spun down for 15 minutes at 4°C at 22,000g and 10μl of the sheared soluble chromatin solution was used in order to check the shearing efficiency and the rest was kept at 4°C. 5% of each sample was kept as an input while the rest of the supernatants were diluted 5 times with dilution buffer (0.01% SDS, 1.1% triton,1.2mM EDTA,16.7mM Tris pH8, 167mM NaCl) and incubated with 3μg H3K27ac antibody (ab4729) O/N under agitation at 4°C. Next day, protein G-Dynabeads were pre-washed 3 times in ice cold 0,01% Tween-20/1xPBS, pre-blocked for 30 minutes at 4°C with 1% BSA/1xPBS and finally added to each sample (30ul Dynabeads per sample) and incubated for 3.5 hours at 4°C in order to bind the specific chromatin-antibody complexes. Upon IP, beads were washed twice in low salt buffer (0.1% SDS,1% triton, 2mM EDTA, 150mM NaCl, 20mM Tris pH8), twice in high salt buffer (0.1% SDS,1% triton, 2mM EDTA, 500mM NaCl, 20mM Tris pH8), twice in LiCl buffer (0.25M LiCl, 1% NP40, 1% <u>deoxycholic acid</u>, 1mM EDTA, 10mM Tris pH8) and once in TE buffer. DNA was then eluted from the beads by incubating with 150ul elution buffer (1% SDS, 100mM NaHCO3) for 30 minutes at 65°C (vortexing every 10min). Input and bound fractions of supernatants were reversed overnight at 65°C with 20mg/ml proteinase K. Next day samples were treated with 100mg/ml RNase and DNA was purified using a ZYMO Kit (D4014) following manufacturer’s instructions. Finally, 25ng of immunoprecipitated material and input were used for ChIP-seq library preparation using the KAPA Hyper prep kit (KK8502) according to manufacturer’s instructions. Libraries were sequenced on an Illumina NextSeq2000 platform on SR100 mode at the Weill Cornell Μedicine Genomics Core Facility. ChIP-seq data have been deposited in the Short Read Archive (SRA) under the accession codes GSE212992.

### ATAC-seq

ATAC-seq was carried out as previously described with minor modifications^136^. For each cell line 2 replicates were performed and analyzed. Briefly, a total of 50,000 cells were washed with 50 μL of cold 1xPBS and then nuclei were isolated in 50 μL lysis buffer (10 mM Tris-HCl pH 7.4, 10 mM NaCl, 3 mM MgCl2, 0.2% (v/v) IGEPAL CA-630). Nuclei were then centrifuged for 10min at 800g at 4°C, followed by the addition of 50 μL transposition reaction mix (25 μL TD buffer, 2.5 μL Tn5 transposase and 22.5 μL ddH_2_O) using reagents from the Nextera <u>DNA library</u> Preparation Kit (Illumina #FC-121-103). Samples were then incubated at 37°C for 30min. DNA was isolated using a ZYMO Kit (D4014). ATAC-seq libraries were prepared using NEBNext High-Fidelity 2X PCR Master Mix (NEB, #M0541), a uniquely barcoded primer per sample, and a universal primer. Samples were first subjected to 5 cycles of initial amplification. To determine the suitable number of cycles required for the second round of PCR (to minimize PCR bias) the library was assessed by quantitative PCR^136^. Briefly, a 5 μL aliquot of the initial amplification sample was used for 20 cycles of qPCR. Linear Rn versus cycle was plotted to determine cycle number corresponding to 1/3 of maximum fluorescent intensity. For each sample, the remaining 45 μL of initial tagmented PCR product was further amplified for 5 more cycles using Nextera primers. Samples were subject to a dual size selection (0.55x–1.5x) using SPRIselect beads (Beckman Coulter, B23317). Fragment distribution of libraries was assessed with an Agilent Bioanalyzer and finally, the ATAC libraries were sequenced on an Illumina Hi-Seq (2500) platform for 50bp paired-end reads.

### In situ Hi-C

The protocol was performed as previously described^42, 137^ with minor modifications. Hi-C was performed starting with 2 million cells per replicate and using the Arima-Hi-C kit (Arima, A510008) according to manufacturer’s instructions. Approximately 500ng of DNA was used for each Hi-C sample to prepare libraries using the KAPA Hyper Prep Kit (KAPA, KK8502) and performing 5 cycles of amplification. Libraries were sequenced using the Illumina Nextseq 2000 in PE50 mode at Weill Cornell Μedicine Genomics Core Facility.

### In situ 4C-seq

The protocol was performed as previously described with minor modifications^138^. Briefly, 10 million cultured ESC, TSC and XEN cells were fixed with 12 ml of 1% formaldehyde (Thermo Scientific, 28908) in 10% FBS for 10 min at room temperature (RT) (tumbling). Quenching of the cross-linking was performed with the addition of 1.8 ml of freshly prepared ice-cold 1 M glycine (Sigma-Aldrich #500046). Tubes were transferred directly on ice and centrifuged for 5 min 500*g* at 4°C. Cells were washed with 1xPBS and centrifuged for 5 min 500g at 4°C, and pellets were frozen in liquid nitrogen and stored at −80°C. Next, cells were then vigorously resuspended in 1 ml of fresh ice-cold lysis buffer (10 mM tris (pH 8), 10 mM NaCl, 0.2% NP-40, and 1 tablet of complete protease inhibitor (Roche, 04693159001)], transferred to 9 ml of prechilled lysis buffer, and incubated for 20 min on ice. Following centrifugation at 500g for 5 min at 4°C, the pellet was resuspended in 50uL of 0.5% SDS and incubated for 10 min at 65°C. SDS was quenched with 145uL ddH_2_O and 25uL of 10% Triton X-100 for 15 mins at 37°C. At this point, 5 μl of the sample was taken as the “undigested control”. Next, 25ul of CutSmart buffer (NEB, B7204S) was added with 10μl DpnII enzyme (NEB, R0543M) and the samples were incubated overnight at 37°C under agitation (750rpm). Upon first digestion, 5μl of the sample was taken as the “digested control” while the efficiency of chromatin digestion was verified after DNA extraction from 5I of undigested and digested controls and loading in a 1.5% agarose gel. After verification of chromatin digestion (smear between 0.2 and 2 kb), DpnII was deactivated by 20 min incubation at 62°C (under agitation 750 rpm). Ligation of DNA ends between the cross-linked DNA fragments was performed by diluting the samples in 669 μL ddH_2_O and adding 120 μL T4 ligation buffer (NEB, B0202), 60 μL 10mM ATP (NEB, P0756S), 120 μL 10% Triton X-100, 6 μL 20mg/ml BSA and 5 μL 400U/μl T4 DNA Ligase (NEB, M0202) overnight at 16°C (tumbling) followed by 30min at RT. 10μl of the ligated sample was tested as “ligated control,” on a 1.5% agarose gel. The samples were then treated with proteinase K and reverse crosslinked overnight at 65°C. Following RNase treatment, phenol/chloroform extraction and DNA precipitation, the pellets were dissolved in 100 μL of 10mM Tris pH 8 and incubated for 1 hour at 37°C. Efficiency of extraction and purification were verified on a 1.5% agarose gel. For the second digestion 20 μL of 10x buffer B (Fermentas), 10 μL Csp6I (Fermentas, ER0211), 80 μL ddH_2_O were added to the DpnII-ligated 3C template and samples were incubated overnight at 37°C under agitation (750rpm). Csp6I was inactivated at 65°C for 20 min, and DNA fragmentation was tested on 1.5% agarose gel. A second ligation was performed by adding 300 μL T4 ligation buffer, 150 μL 10mM ATP, 5μL T4 DNA Ligase, and ddH_2_O to 3mL and incubating overnight at 16°C. After 30 min of incubation at RT, samples were PCI-extracted, ethanol-precipitated, resuspended in 200 μl of sterile water, and purified using the Qiaquick PCR Purification Kit (Qiagen). DNA concentration of each digested sample was calculated using the Qubit brDNA HS assay kit (Invitrogen). For library preparation, primers were designed either around the enhancer or the promoter of lineage specific genes. Library preparation was then performed using the <u>inverse PCR</u> strategy. Briefly, 4×200 ng of 4C-template DNA was used to PCR amplify the libraries using the Roche Expand long template PCR system (Roche, 11681842001) with the following PCR conditions: 94 °C for 2 min, 16 cycles: 94 °C for 10 seconds; [primer specific] °C for 1 min; 68 °C for 3 min, followed by a final step of 68 °C for 5 min. Amplified material was pooled, and primers were removed using SPRIselect beads (Beckman Coulter, B23317). A second round of PCR with the following conditions: (94 °C for 2 min, 20 cycles: 94 °C for 10 seconds; 60 °C for 1 min; 68 °C for 3 min and 68 °C for 5 min) was performed using the initial PCR library as a template, with overlapping primers to add the P5/P7 sequencing primers and indexes. The samples quantity and purity were determined using a NanoDrop spectrophotometer while the 4C PCR library efficiency and the absence of primer dimers were reconfirmed by Agilent Bioanalyzer. For each cell line 3 replicates were performed, and the libraries were sequenced on a HiSeq4000 in SE150 mode at Weill Cornell Medicine Genomics Core Facility. All the 4C-seq primer sequences are provided in Supplementary Table 7.

### H3K27ac HiChIP

ESC cells were processed for each HiChIP replicate using the Abcam H3K27ac antibody (ab4729) and following the HiChIP protocol as previously described^42^. TSC, XEN and EpiSC cells were used for each HiChIP replicate using the Arima-HiC+ kit (Arima, A101020) and the H3K27ac antibody (active motif H3K27ac 91193) according to manufacturer’s instructions with few modifications. The efficiencies of H3K27ac antibodies were tested by ChIP-seq, and both antibodies resulted in similar distribution and number of peaks. In order to improve the sonication efficiency, a modified lysis buffer was used containing 10mM Tris pH8, 1mM EDTA and 0.5% SDS. Prior to over-night incubation with the antibody the sample was diluted in a buffer to bring it back the original composition of the Arima R1 buffer (10mM Tris pH8, 140mM NaCl, 1mM EDTA, 1% triton, 0.1% SDS, 0.1% sodium deoxycholate). 5ng of immunoprecipitated DNA material was used to make libraries using the Swift Biosciences Accel -NGS 2S Plus DNA Library Kit (Cat #21024) according to manufacturer’s instructions and performing between 8-14 cycles of amplification for all samples. Final libraries were sequenced using the Illumina Nextseq 2000 in PE50 mode.

### ChIP-seq analysis

All single-end sequenced reads were aligned to mouse genome (mm10) with Bowtie2 (version 2.3.4.1) ^139^ and “--local –very-sensitive-local” option. Samtools, “MarkDuplicates” from picard tools and bedtools were used to filter out low quality reads (MAPQ<20), duplicate reads, chrM and blacklisted regions. Filtered reads were used to call ‘broad’ peaks with MACS2 (version 2.1.1) and default settings. Non overlapping peaks from replicates were filtered out and only common peaks were used. Identification of the 5 enhancer groups was performed with K-mean clustering on the enhancer atlas of all H3K27ac peaks in the 3 cell lines under investigation. The same pipeline was used for all published ChIP-seq datasets that were included in this study.

### ChIP-exo analysis

All paired-end sequenced reads were aligned to mouse genome (mm10) with Bowtie2 (version 2.3.4.1) ^139^ and “--local –very-sensitive-local” option. Reads were trimmed to 36bp and we used samtools, “MarkDuplicates” from picard tools and bedtools to filter out low quality reads (MAPQ<20), duplicate reads, chrM and blacklisted regions. Filtered reads were used to call ‘broad’ peaks with MACS2 (version 2.1.1) and default settings. Non overlapping peaks from replicates were filtered out and only common peaks were used.

### ATAC-seq analysis

All paired-end sequenced reads were aligned to mouse genome (mm10) with Bowtie2 (version 2.3.4.1) ^139^ and “--local –very-sensitive-local -I 10 × 2000” option. Samtools, “MarkDuplicates” from picard tools and bedtools were used to filter out low quality reads (MAPQ<20), duplicate reads, chrM and blacklisted regions. All filtered reads were corrected for Tn5 insertion at each read end by shifting +4/-5 bp from the positive and negative strand respectively. MACS2 with ‘--broad’ option and default settings were used to call peaks. Non overlapping peaks from replicates were filtered out and only common peaks were used. Peak center (summit file) generated with MACS2 with ‘--narrow’ option was extended to 100bp (+/-50bp) for motif search and all overlapping summits were merged to form an accessibility atlas which was used as background for motif and ChIP enrichment with LOLA R package.

### RNA-seq analysis

Tophat2 (version 2.1.1) with default setting and “-r 200 –mate-std-dev 100” was used to align paired-end sequenced reads to mouse genome (mm10). Sorting of aligned reads was performed with samtools and reads were assigned to protein coding and long-non coding genes (Mus_musculus.GRCm38.95.gtf) with the use of htseq-count^140^ and ‘-m intersection-nonempty’ option. Identification of differential expressed genes was performed with DESeq R package and p-adj <0.01 and fold change 2 as cut offs. All expressed genes significantly upregulated in the respective cell line compared to the other 2 lineages (TPM>1, LogFC >2 and p-adjusted <0.01) were considered lineage specific genes. Housekeeping genes used for analysis were downloaded from HRT Atlas v1.0 database (PMID: 32663312).

### Hi-C analysis

Hi-C data were pre-processed using HiC-bench platform^141^. Read pairs with low MAPQ, self-ligated fragments and short-range interaction (<40kb) were filtered out prior to downstream analysis. ICE normalized matrices at various resolutions and .hic files were generated with both Hi-C-bench and juicer tools ‘pre’ option^142^. Compartment analysis was performed at 100kb resolution with the use of CscoreTool (version 1.1)^143^ for each experiment and chromosome separately with ‘minDis 1000000’ option and 100kb bins. Compartments were assigned to ‘A’ (active) and ‘B’ (inactive) based on gene density for all bins. Topologically associated domains, boundaries and insulation scores were calculated with Hi-C-bench pipeline by using the ‘domain’ operation on the Hi-C matrix at 40kb resolution. Aggregate peak analysis (APA) plot was generated with APA package from juicer tools (version #1.22.01) and ‘-w 10 -r 5000’ settings.

### TAD identification

The HiC-Ratio algorithm integrated in HiC-Bench with default parameters, which computes insulation scores as described in^141^.

### IntraTAD activity analysis

Iteratively corrected matrices were re-normalized by dividing each bin value by the sum of all the values in the same distance bin in the same chromosome (distance normalization), or by the total number of valid pairs (‘cpm’)^144, 145^. All the TADs identified in the control sample were used as the reference TADs to compute the intra-TAD activity changes. The set of reference TADs between the 2 samples, S1 (control) and S2 (treatment), were denoted as set *T*. A paired two-sided *t*-test was performed on each single interaction bin within each reference TAD between the 2 samples. We also calculated the difference between the average scores of all interaction intensities within such TADs and the TAD interaction log fold change. Finally, a multiple testing correction is performed by calculating the FDR on the total number of TAD pairs tested. The TAD interaction change for each *t* in *T* is calculated as follows:

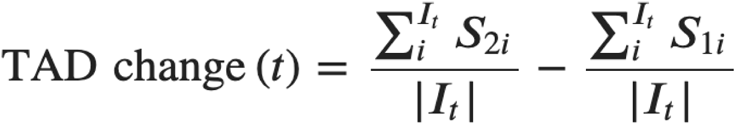

We classified the reference TADs in terms of Loss, Gain or Stable intra-TAD changes by using the following thresholds: FDR < 0.01 and absolute TAD interaction log fold change >0.25, absolute TAD interaction change >0.1.

### Connectivity Analysis

We used the ‘boundary-scores’ pipeline in HiC-Bench with ‘connectivity’ parameters. For each genomic bin (40k or 100k) it computes the sum of the ic-normalized interactions in a distance of 0.5 Mb or 2 Mb.

### 4C-seq analysis

Demultiplexing, trimming of VP and resizing of sequence reads to 35 bp was performed with fastq_trimmer while fastx_clipper (Fastx-toolkit version 0.0.14) was used for selecting reads with RE site next to the VP. Alignment of sequence reads was performed with Bowtie2 (version 2.3.4.1)^139^ and “--local –very-sensitive-local” option to mouse genome (mm10 genome version). Both Samtools (version 1.7-2)^146^ and Bedtools (version 2.26.0)^147^ were used for filtering low quality reads (MAPQ<20), chrM and blacklisted regions. All reads were assigned to an RE site and all reads within 2Mb of the VP excluding the first 2 REs in both sides of the VP were used for CPM normalization. BigWig were generated with bedtools genomecov and bedGraphToBigWig^148^. All regions between RE sites were assigned RE normalized CPM value of the first RE while rolling mean of 21 RE was performed in R (version 4.0.4) with “rollmeanr”.

### HiChIP analysis

All sequencing files were processed with HiC-Pro pipeline (version 3.0.0). Bed files with *in silico* digestion of the mm10 genome by MboI or Arima restriction enzymes were generated with ‘digest_genome.py’ tool from HiC-Pro and were used for assigning mapped reads to DNA fragments. Valid deduplicated reads from replicates were merged and were used for loop calling at 5kb resolution with FitHiChIP (release 9.0) and coverage bias regression option active. Loops with one peak in either of the 2 interacting regions (IntType = 3: “peak to all” option), sizes between 10-2000kb and FDR <0.01 for ESC, TSC, XEN and <0.05 for EPISC were considered valid.

Loops identified by FitHiChIP were separated into 5 categories (Promoter-Promoter, Promoter-Enhancer, Enhancer-Enhancer, Promoter-X, Enhancer-X) based on the presence of an Enhancer or a TSS within their 5kb anchors. Each anchor containing a TSS was characterized as a Promoter anchor (P) and presence of a H3K27ac peaks in regions with no TSS were characterized as Enhancer anchors (E). Lack of both marks (P and E) resulted in identification of X-anchors. Multiconnected anchors were considered to be hubs and based on the type of the multi-connected anchor they were separated into Promoter, Enhancer and X – hubs.

### Gene Ontology

Gene ontology of genomic regions in bed format was performed with GREAT (version 4.0.4) for mm10 genome and ‘Basal plus extension’ with ‘plus Distal’ option extended to 50kb. GO terms from biological processes with p-value <0.01 were scored as significant. Gene ontology of genes was performed in R with goprofiler2 with the use official gene symbols by setting user_threshold to 0.05 and “g_SCS” as correction method. KEGG, GO:BP and WP sources were selected for gene annotation. Additional Gene Ontology was performed by metascape online analysis. Default options were chosen including terms from Wikipathways, Reactome Gene sets, KEGG pathways and GO biological processes.

### Super-enhancer analysis

ROSE pipeline was used to call super-enhancers in all cell lines. For each cell line .bam files of both replicates were merged and converted to GFF according to ROSE pipeline and H3K27ac common for each cell lines were used for super-enhancer identification. Enhancer regions at 12.5kb distance were stitched into one with ‘-s 12500’ option active.

### Region Enrichment

LOLA (version 1.8.0)^46^ software in R was used to calculate enrichment of ChIP-seq data and TF motifs on mouse genome. LOLA database was expanded based on available published ChIP-seq data for ESC, TSC and XEN^42^. Overlap and enrichment of accessible sites, super-enhancers and H3K27ac peaks with mm10 LOLA region database was performed by comparing accessible sites overlapping regions of interest with all accessible regions as control. In addition to ChiP-seq enrichment we generated a database that contained 726 motifs in bed format as extracted from PWMScan database^149^ for “JASPAR CORE 2020 vertebrates” and “HOCOMOCO v11 Mouse TF Collection” motifs. Significant enrichment of transcription factors and motifs was scored based on p-value levels (<10^(−3)).

### Modeling

Random Forest methodology was used for classification of gene expression levels and gene expression level prediction. A set of 28 variables that contain information from 1D (H3K27ac, ATAC-seq) and 3D (HiChIP) experiments were calculated for all hubs in our 4 cell types (Supplementary Table 6). After eliminating features with high correlation among them from 1D, 3D and combined 3D we ended up with 10 features. Recursive feature elimination (rfe function in “caret” library in R) was used for feature selection which led to the use of 8 out of the 10 features both in classification and regression Random Forest models.

Classification of hubbed genes based on their expression levels was achieved by separating looped genes into 10 equally sized groups (Q1 to 10). Cross validation was performed with “leave one chromosome out” method (L.O.C.O.) where we train our data in all chromosomes but one which we use for testing. This process is repeated until we leave every chromosome out of the training test for chromosomes 1-19 and chrX. AUC and correlation scores are calculated in each round of LOCO (n=20) and average AUC and correlation is calculated for all of our models tested (promoter, linear 2D and 3D). TSC promoter hubs for Q1 and Q10 were used for training, with ntree=1000 and mtry=floor (sqrt(# Variables) in TSC and tested classification of Q10 and Q1 gene groups in ESC, XEN and EPISC. In order to evaluate the models, we calculated average AUC score for each model in all cell lines. None of the model showed over-fitting since both training and testing sample showed similar accuracy. The same methodology was used to identify differential expression. For each cell type pair (n=6) we merged looped genes and calculated the difference for all of our 8 variables. We selected TSC/EPISC pair as our initial dataset which was split into training and test dataset as before with LOCO by selecting the Q1 and Q10 promoter hubs based on fold change. Random Forest was applied as before and average AUC scores were calculated for the rest cell type pairs (n=5, ESC/XEN, ESC/EPISC, TSC/ESC, TSC/XEN, XEN/EPISC).

Gene expression prediction was achieved with Random Forest regression model and ntree=1000 and mtry=floor(#Variables/3). Again, TSC was used for training and testing for all hubbed genes. The same steps were followed when we performed RF to predict fold changes between cell type pairs. Evaluation of RF model was performed with average spearman rank correlation coefficient.

To estimate the effect of each enhancer in our cell lines we performed *in silico* perturbation of each hub by removing one enhancer at a time in ESC, XEN and TSC. All 8 variables (hub metrics) were recalculated after each enhancer removal and gene expression levels were estimated based on the new hub metrics. In-silico perturbation was estimated as the percentage of change between Predicted and In-silico predicted gene expression levels for each of the genes and were separated into two groups based on their gene expression changes (Perturbed vs Not perturbed).

### Hi-ChAT score calculation

HiChAT score is calculated for each promoter anchor taking into account accessibility, enhancer and loop strength similar to ABC score^46^. For each gene only their interacting-looped enhancers within a 4Mb regions were used. ATAC signal was used for estimating accessibility of the enhancer identified by H3K27ac. For each promoter hub HiChAT was calculated with the following formula:

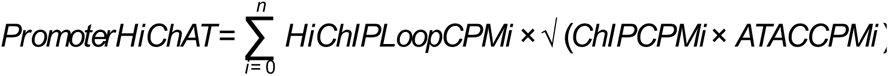

where n is the number of connected enhancer anchors for a given promoter. HiChAT calculation provides an ABC-like score^46^ for all promoters by aggregating the Activity by contact signal of all connected enhancers. Two HiChAT scores (1 & 2) were generated by calculating the combined ATAC/H3K27ac signal at the enhancer and accessible regions respectively and tested in our gene expression predicting models.

### Virtual 4C

Valid paired end reads from Hi-C and HiChIP were used to generate bigwig files representing the cis-interactions for selected genes and enhancers. We generated successive windows at 5kb resolution overlapping by 90%. After isolating paired end reads that overlap with the bin or interest (TSS or enhancer center) we count the number of reads in all overlapping windows at 2Mb distance from the region of interest. Read counts are normalized to the total number of reads within this 2Mb after removing all reads that overlap with the region of interest for each cell type and bedGraphs generated are converted to bigWig files with the use of kent-tools^148^.

### Statistical methods and plots

Median comparisons were performed with the use of two-sided Wilcoxon rank test in R while two-sided Fisher’s exact test was used to compare enrichment or differences in distribution. Student’s T-test was used to compare C-score and insulation levels between different cell lines. In any of the above methods significance was estimated based on p-value levels (<0.05). All heatmaps, barplots, enrichment dot plots, scatter plots, boxplots and ROC curves were generated in R. K mean-heatmap of H3K27ac signal enrichment was generated with DeepTools^150^ and bigWigCompare tool. All genome data visualization were generated in IGV browser with the use of bed files for genomic regions, big wigs for signal enrichment and ARCs for loops.

## Data availability

All genomic datasets generated in this study (ChIP-seq, ATAC-seq, RNA-seq, 4C-seq, HiC and HiChIP) have been uploaded in the Gene Expression Omnibus (GEO) under GSE213645 accession number. Source data are provided with this paper.

## Code availability

Custom R scripts used for data analysis in this study have been developed in our lab and are available upon request.

## ACKNOWLEDGEMENTS

We are grateful to all member from the Apostolou and Stadtfeld groups for critical reading of the manuscript and input on this work along the way. We also thank Julian Pulecio and Danwei Huangfu for advice on the functional experiments and Christina Leslie for advice on the modeling. We also thank the Weill Cornell Μedicine Genomics Core Facility and the Flow Cytometry Core Facility. This work was partly supported by a HiChIP research grant from Arima Genomics. DM was supported by the T32 HD060600. EA is a recipient of the Mark Foundation Emerging Leader Award and supported by the NIH (1R01GM138635, 1U01DK128852, RM1GM139738) and the Tri-Institutional Stem Cell Initiative of the Starr Foundation.

## AUTHOR CONTRIBUTIONS

EA and AP conceived and designed the study and analyses with input from DM, ES, MS, AKH and AT. DM and ES performed the genomic and all the functional experiments. DCG assisted with genomic experiments. VG provided help with TSC and XEN cell lines, while LE provided material for the EpiSC genomics experiments. CU assisted with HiChIP visualization. UL assisted with CTCF ChIP-exo in ESC. AP performed all computational analyses with help from JRH, AK and guidance from AT and EA. EA wrote the manuscript together with DM, ES and AP and input from all authors.

## Extended Data

**Extended Data Fig. 1. Related to Figure 1.**
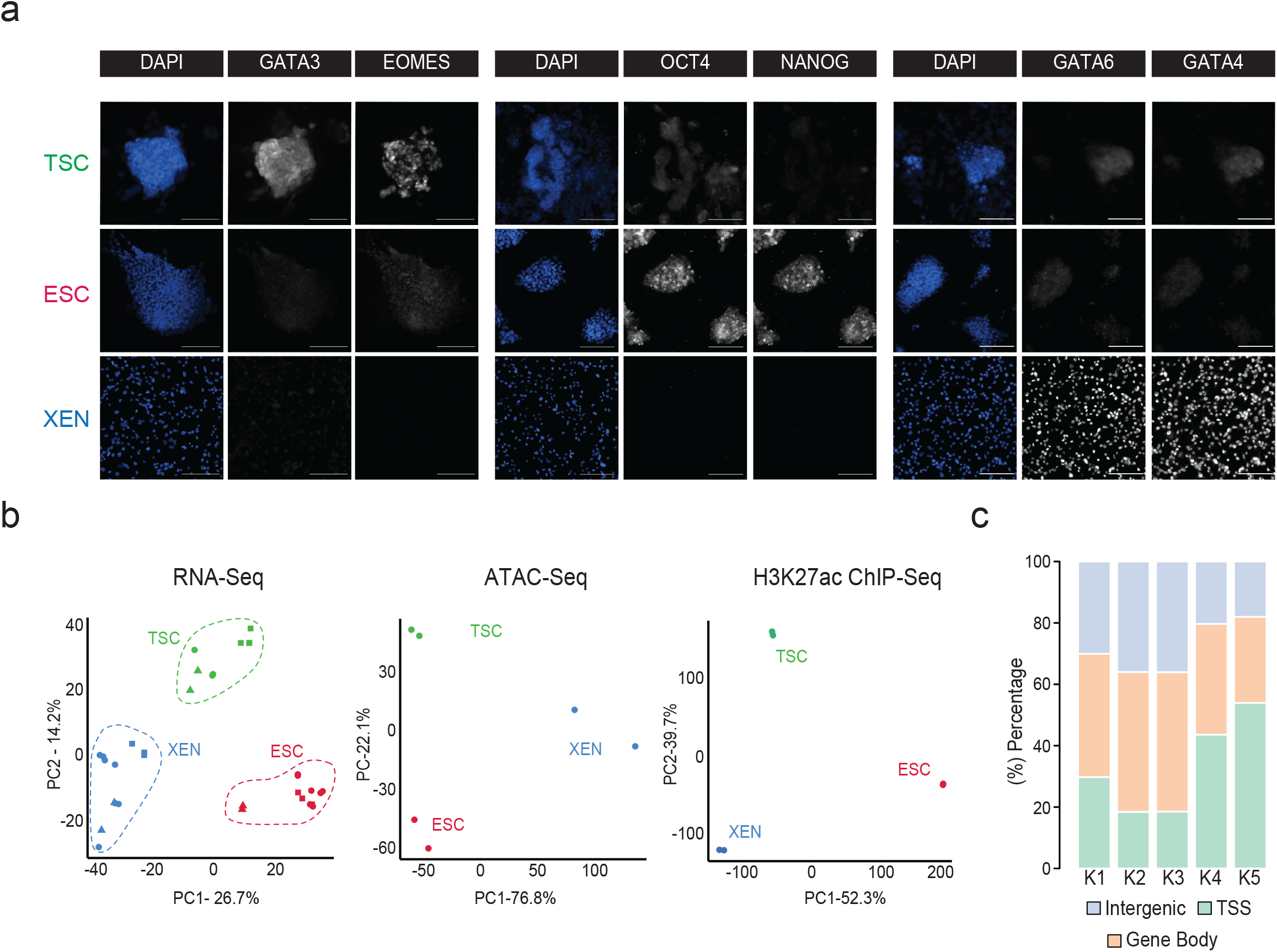
a. Representative single (xy) stack epifluorescence images of immunofluorescence experiments showing expression of key lineage markers (greyscale) in TSC, ESC and XEN cells. Cells were counterstained with DAPI (blue) for DNA content. Scale bar 100μm. b. Principal component analysis (PCA) of all TSC, ESC and XEN replicates based on their RNA-seq, ATAC-seq and H3K27ac ChIP-seq profiles. PCA plots were designed based on the top10% of most variable genes or peaks in all three cell lines. In each plot, circles indicate the experimental data presented in this study, while squares and triangles correspond to publicly available RNA-seq data (Supplementary Table 8) or independent -unpublished-studies from our lab, respectively. c. Stacked barplot showing the distribution of H3K27 occupancy among intergenic regions, gene bodies or TSS (promoter +/-1.5kb) for each K-Mean cluster as identified in Fig.1c. *Note:* all statistics are provided in Supplementary Table 9.

**Extended Data Fig. 2. Related to Figure 2.**
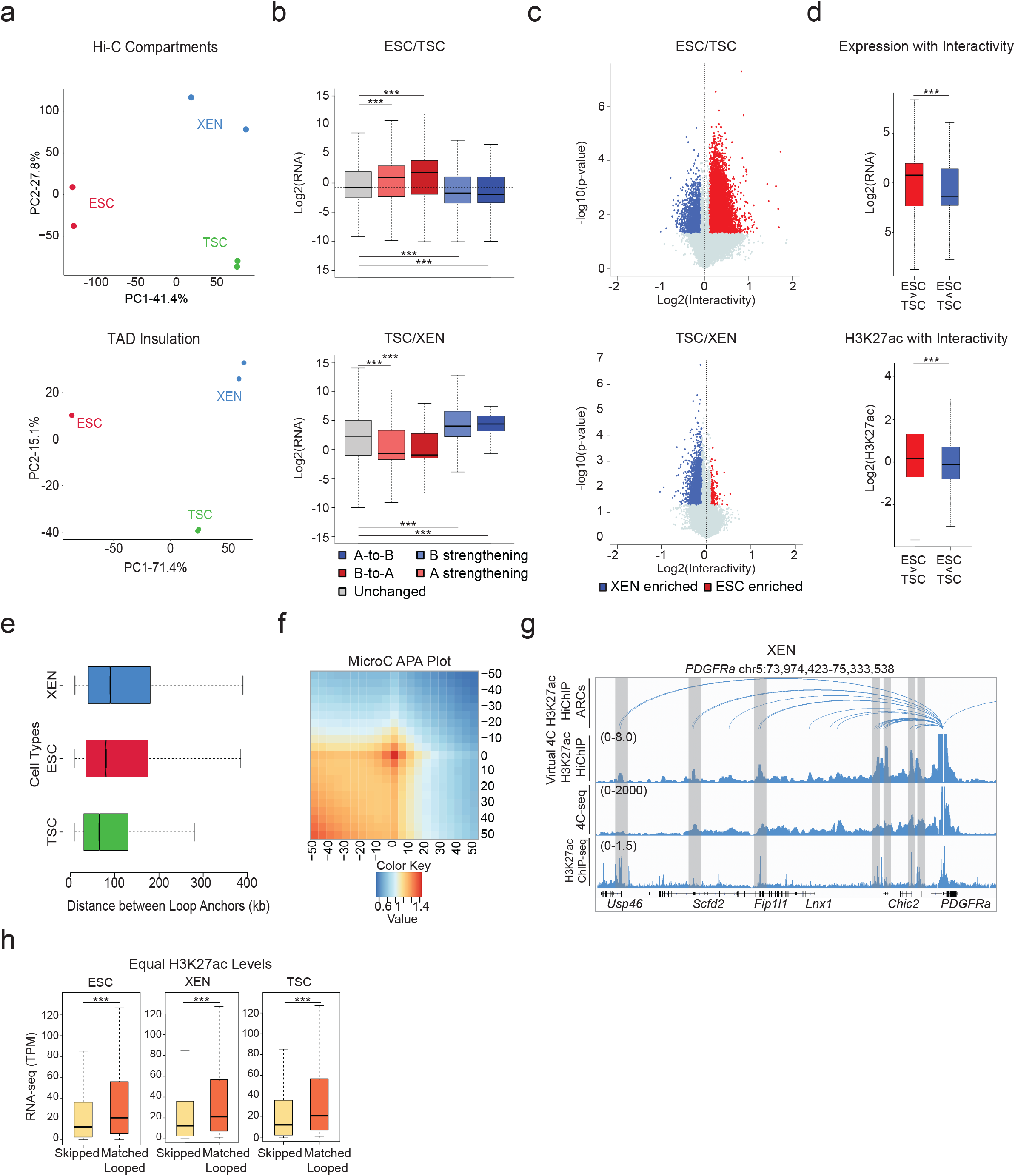
a. Principal Component Analysis (PCA) plot of all lineages and replicates based on their compartment scores at 100kb resolution (top) and on their TAD insulation levels at 40kb resolution (bottom). b. Boxplots showing median expression changes between ESC and TSC, and TSC and XEN cells of genes that reside either in unaltered compartments (grey box and dashed line) or compartments that undergo shifts and changes as described in Fig 2b. c. Volcano plot showing differential Hi-C interactivity at 40kb resolution between ESC – TSC and TSC - XEN. X-axis shows the difference of the interactivity levels while y-axis shows -log10(p-value) as calculated by two-sided Student’s t-test. Significant changes (p-value<0.05 and Diff>0.1 or <-0.1) are noted with blue and red color. d. Boxplots showing gene expression and enhancer strength changes between ESC-TSC regions that underwent connectivity changes as described in (Fig. 2c). e. Boxplot comparing the sizes of HiChIP-detected loops in the three cell lineages. f. Aggregate peak analysis (APA) showing the aggregate signal of MicroC data in ESC ^39^ centered around ESC HiChIP interacting regions as identified by FitHiChIP2.0 at 5 kb resolution. APA score is calculated as the ratio of the number of contacts of MicroC interacting regions (center bin) to the mean numbers of contacts in the lower left corner. For further details see also Supplementary Table 4. g. IGV tracks aligning H3K27ac HiChIP results (arcs on top and virtual 4C representation in the middle) with the 4C-seq normalized signals around *PDGFRA* promoter in XEN along with corresponding H3K27ac ChIP-seq occupancy. Interactions detected by both HiChIP and 4C-seq are highlighted. The average 4C-seq signals and the H3K27ac ChIP-seq were normalized to the sequencing depth derived from two biological replicates. h. Boxplot showing the median expression levels of a curated list of skipped and looped genes in ESC, XEN and TSC. These genes were selected to have similar range of H3K27ac signal around their promoters. Asterisks indicate significance (p-value<0.05), as calculated by Wilcoxon rank sum test. For further details see also Supplementary Table 4. *Note:* all statistics are provided in Supplementary Table 9.

**Extended Data Fig.3. Related to Figure 3.**
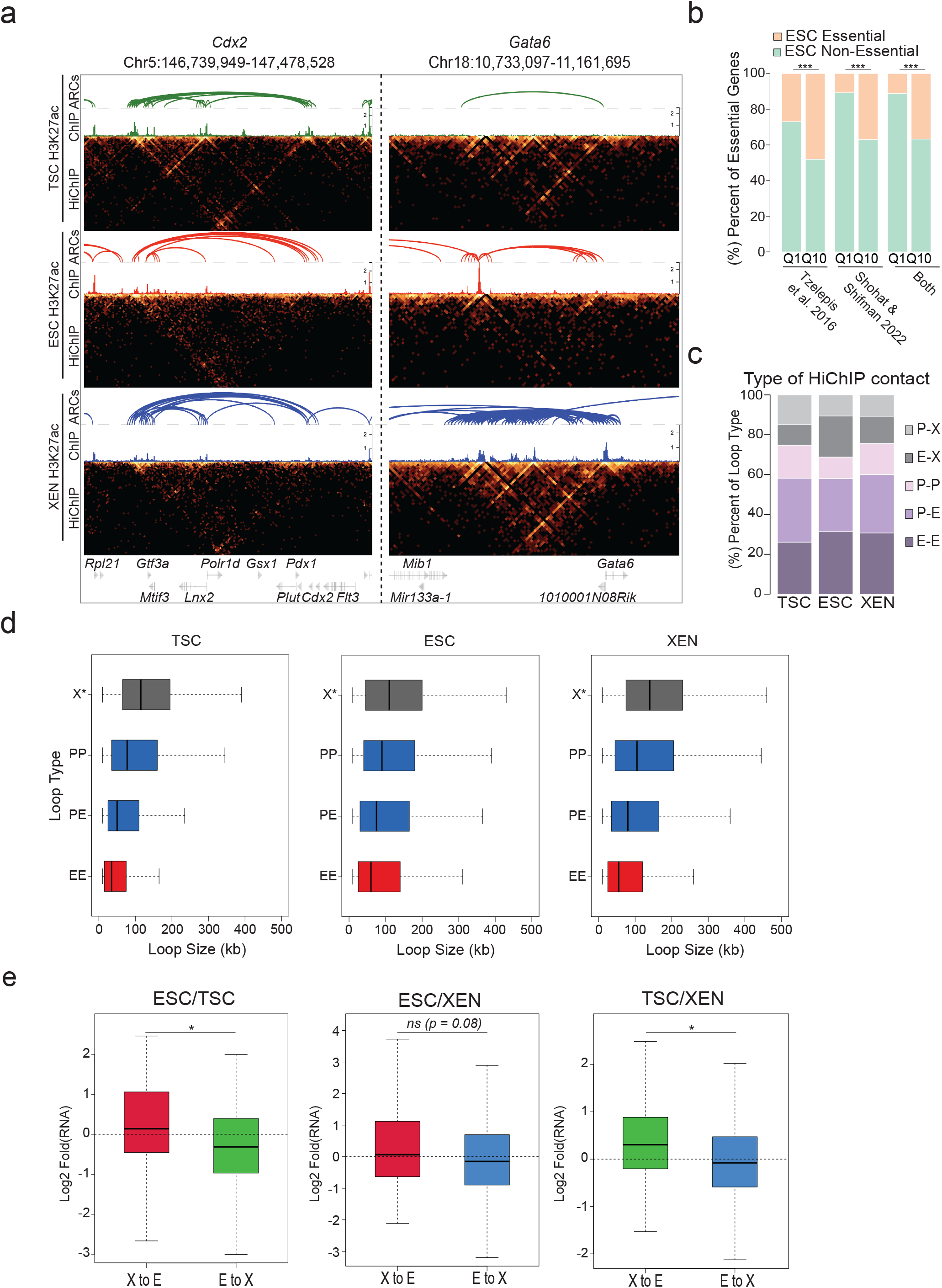
a. HiGlass visualization of H3K27ac HiChIP results around a TSC related hub (Cdx2) and a XEN-related hub (Gata6) in TSC, ESC, and XEN along with the corresponding H3K27ac HiChIP derived arcs and H3K27ac ChIP-seq signals. Interacting scores are presented in 5kb resolution. b. Barplot showing the percentages of essential genes -as identified in two recent studies ^95, 96^-within the least (Q1) versus most (Q10) connected hubs. The preferential enrichment of essential genes in Q10 is significant (p-value<0.001, Fisher’s exact test). c. Stacked barplots showing the proportions of different HiChIP loop subtypes in TSC, ESC and XEN cells. Loops were separated into 5 chromatin interaction categories based on the presence of regulatory elements, such as promoter/TSS (P) or putative enhancer (E, H3K27ac peak). X-anchors were defined as anchors that do not contain any TSS nor an H3K27ac peak. d. Boxplot showing the size distribution of X loops (X-E and X-P) compared to E-E, E-P and P-P loops in all cell lines. e. Boxplots showing expression changes between any two cell types around multiconnected genes (n>=5 in both cell types of interest), when at least one of their conserved anchors switches chromatin states: either from X-to-E (enhancer gain) or from E-to-X (enhancer loss) Asterisks indicate significance < 0.05. (See also Supplementary Table 9). *Note:* all statistics are provided in Supplementary Table 9.

**Extended data Fig. 4. Related to Figure 4.**
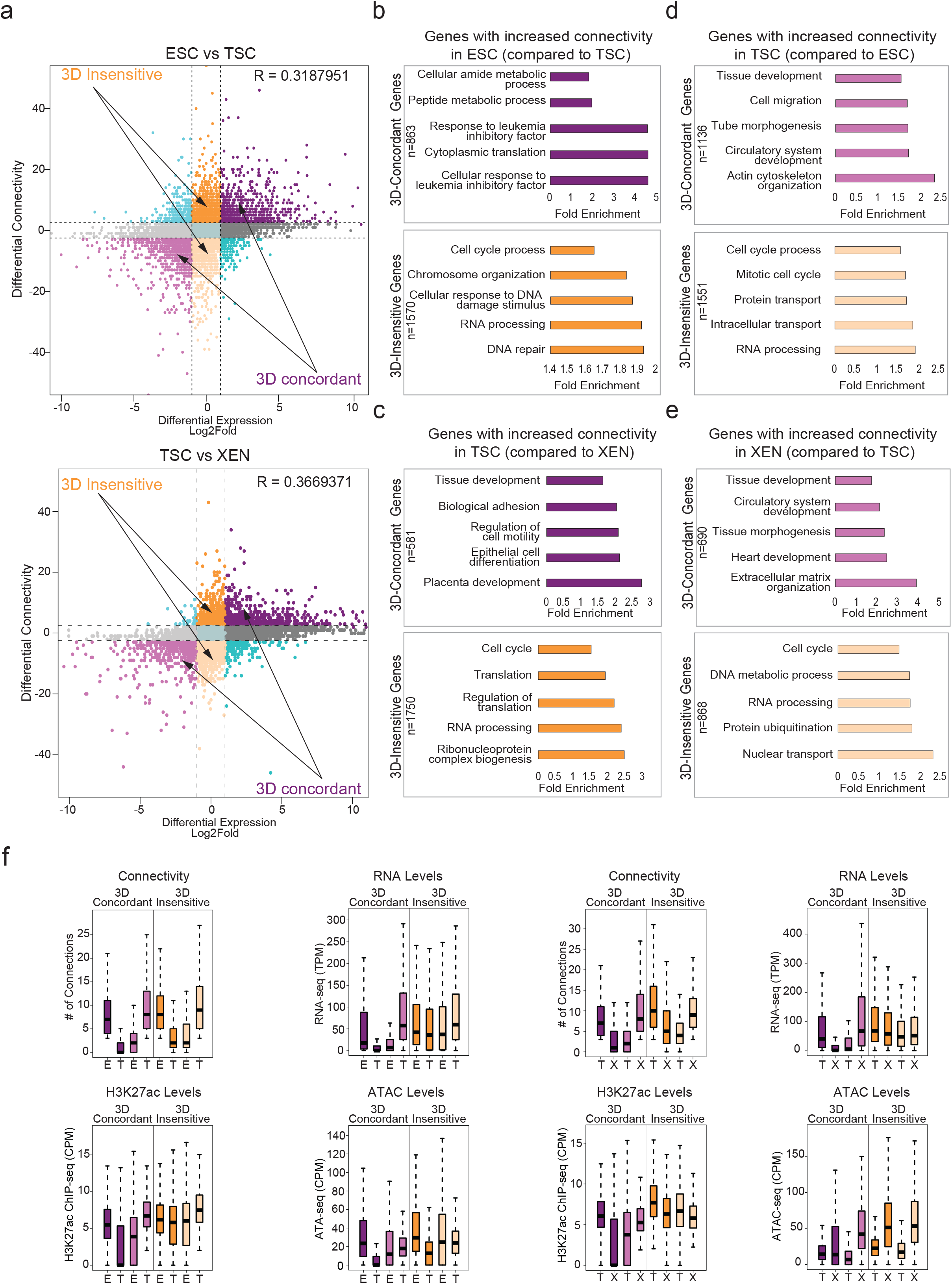
a. Correlation between differential HiChIP connectivity/hubness and differential gene expression in connectivity and differential gene expression in ESC and TSC cells (top) and TSC and XEN cells (bottom). R represents Spearman correlation identifies distinct groups of genes. We focus on the two most prominent groups: 3D-insensitive genes, defined as genes with differential connectivity >3 but no transcriptional changes (log2FC <1 or >-1) and 3D-concordant genes for which connectivity and expression changes (log2FC >1 or <-1) positively correlate (Supplementary Table 5). b-e. Gene ontology analysis depicting the most significant biological processes enriched in the 3D concordant and 3D-insensitive genes in each pairwise comparison (ESC vs TSC and TSC vs XEN) as defined in (a). All genes in A compartments were used as background. For further details see also Supplementary Table 3. f. Comparison of connectivity, gene expression levels as well as H3K27ac and ATAC CPM levels on the promoters of 3D-concordant and 3D-insensitive genes between ESC and TSC cells (left) and TSC and XEN cells (right) as defined in (a). Insensitive genes show higher levels of connectivity, H3K27ac, ATAC and expression in both cell types. Wilcoxon rank sum test was used for all comparisons (Supplementary Table 9). *Note:* all statistics are provided in Supplementary Table 9.

**Extended data Fig. 5. Related to Figure 5.**
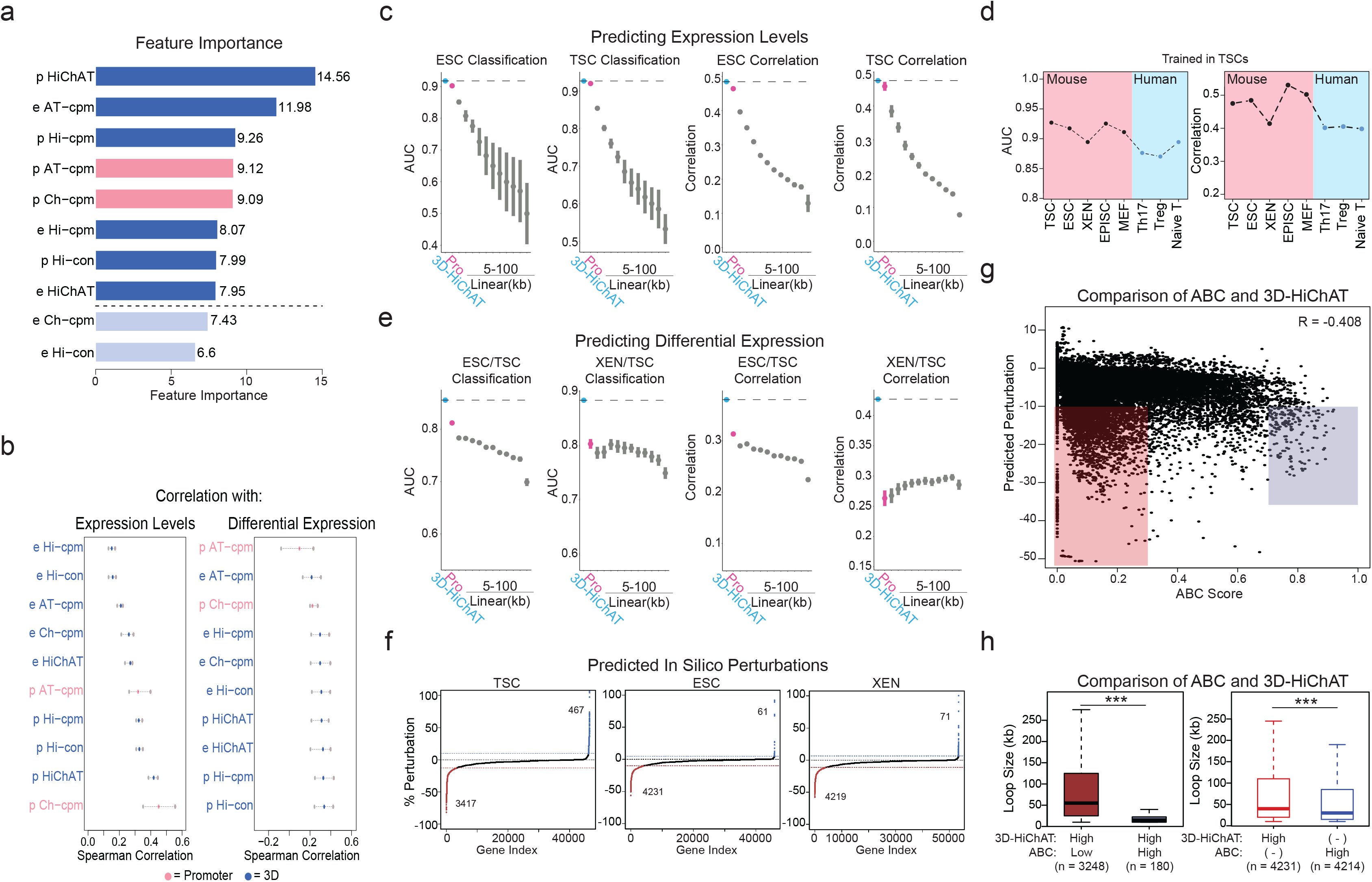
a. Barplot of feature importance scores calculated using recursive feature selection method for predicting gene expression levels, shows that 8 out of 10 features scored as important ranked by high to low. Pink indicates 1D features, while blue indicates 3D variables. Light blue color indicates the features that were not selected for model training. For further details see also Supplementary Table 6. b. Spearman correlation values between each of the 10 variables considered for our 3D model with gene expression levels (left) and differential expression levels (right). For each feature the dots represent the minimum, mean and maximum correlation score from 4 tested cell lines (ESC, TSC, XEN and EPISC) (left) or from 6 differential analysis pairs (right). For further details see also Supplementary Table 6. c. Area Under Curve (AUC) scores and Spearman Correlation scores generated for predicting classification of gene expression (top 10% high vs low expressing genes, left graph) and absolute levels (right graph) in ESC or TSC cells using each of our 3D-HiChAT, Promoter-1D and Linear-1D models across various distances from the TSS (5kb-100kb). Each dot represents the average score across all 20 chromosomes using the LOCO approach, while error bars show standard deviation. See also Extended Data Figure 5 for the rest of the cell lines and comparisons. For further details see also Supplementary Table 6. d. Plots showing AUC and Sperman correlation scores for predicting classification of gene expression (top 10% high vs low expressing genes, left graph) and absolute levels (right graph) using 3D-HiChAT model (Trained in TSCs) in various lineages including mouse lineages: TSCs, ESCs, XEN, EpiSCs and MEFs^42^ and published data from human lineages: Naïve T cells, T-Helper 17 Cells (Th17), and T regulatory cells (Tregs)^151, 152^. e. Area Under Curve (AUC) scores and Spearman Correlation scores generated for predicting differential expression classification (top 10% up or downregulated, left) and fold change expression (right) between XEN and ESCs using each of 3D-HiChAT, Promoter-1D and Linear-1D models across various distances from the TSS (5kb-100kb). Each dot represents the average score across all 20 chromosomes using the LOCO approach, while error bars show standard deviation. For further details see also Supplementary Table 6. f. Ranked perturbation scores (%) as predicted by *in silico* perturbations of ∼46K E-P pairs in ESC, ∼46.7K in TSC and ∼53.1K in XEN using the 3D-HiChAT model. The dotted horizontal lines indicate the selected cut-offs for impactful or not perturbations, defined as the points on the curves where the slope of the tangent is >1 (blue) or <-1 (red). The latter represent putative functional enhancer-promoter pairs, since in silico perturbation of the enahcers results in reduced predicted gene expression levels. g. Scatterplot comparing for each anchor the predicted perturbation scores from our 3D-HiChAT model with the respective ABC scores. The R Spearman correlation value is shown on the top. h. Boxplots showing that enhancers with high 3D-HiChAT-predicted perturbation scores and low ABC scores (red) are significantly more distal to their target genes (loop size) than those with concordant high scores in both models (blue) (left plot). Similarly, comparison of all enhancers/anchors with either high 3D-HiChAT predicted perturbation scores (perturbation <-10%, red) or high ABC scores (ABC>0.7), show that 3D-HiChAT predicts potentially functional enhancers at larger distances (right plot). *n*, indicates the number of anchors used for each comparison. Asterisk indicates significance, p-val<0.001 *Note:* all statistics are provided in Supplementary Table 9.

**Extended Data Fig. 6. Related to Figure 6.**
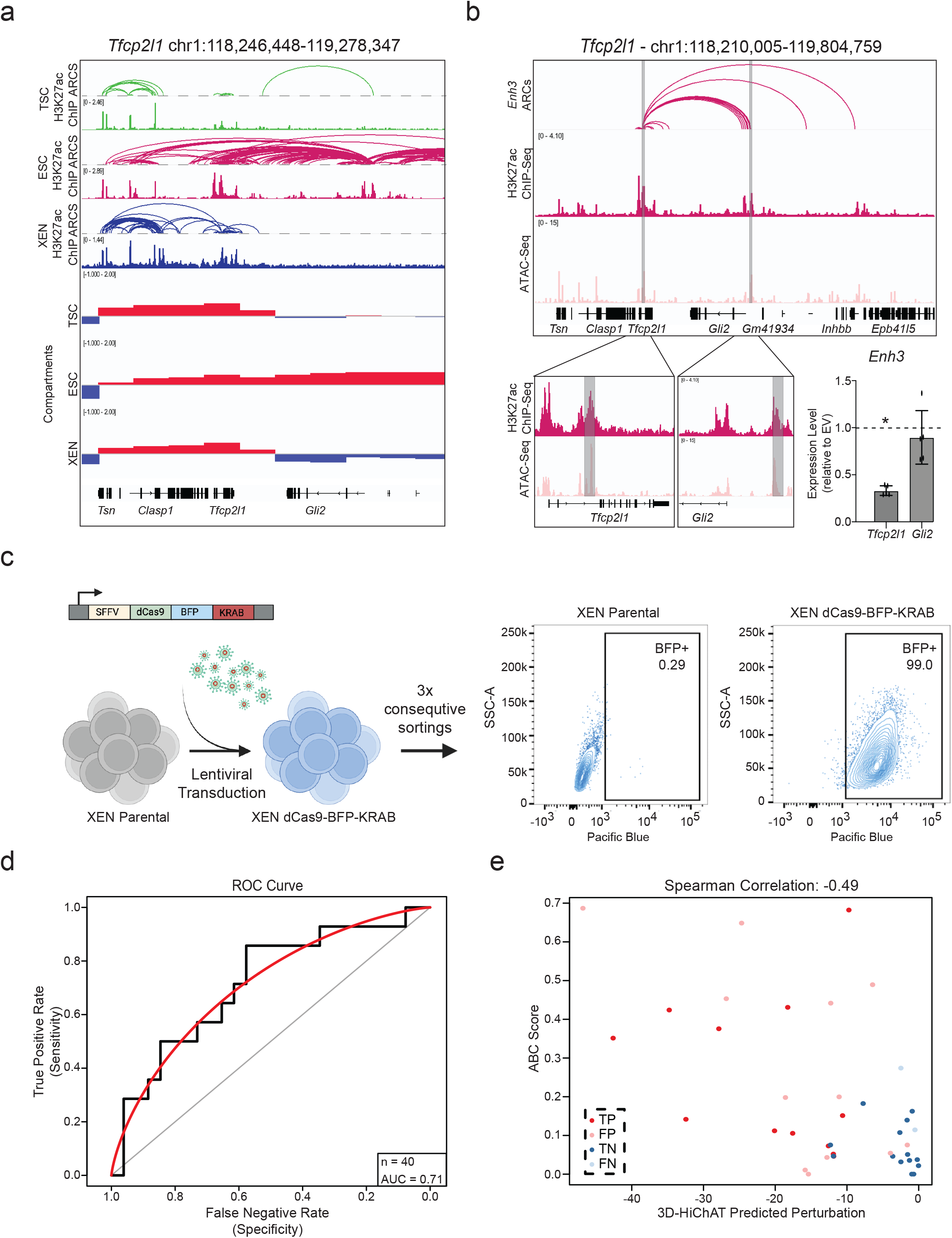
a. Visualization of the *Tfcp2l1* Locus showing H3K27ac HiChIP arcs, H3K27ac ChIP and Compartment c-scores called by HiC for TSC, ESC, and XEN. Notably, a group of putative enhancers upstream of *Gli2* are uniquely expressed and only in an A compartment in ESCs. b. IGV tracks of the *Tfcp2l1-Gli2* locus showing the two enhancers chosen for functional validation, Enhancer 3 and 14. H3K27ac Hi-ChIP derived arcs originating from both enhancers are shown as well. RT-qPCR showing relative expression levels of *Tfcp2l and Gli2* upon CRISPRi perturbation of Enh3 compared to control cells infected with empty vector (EV). Dots indicate independent experiments (n=3). Asterisks indicate significance, with p-value <0.05, as calculated using unpaired one-tailed t-test. c. Schematic showing experimental strategy for generating a stable XEN line expressing dCas-BFP-KRAB (CRISPRi) as shown by the representative FACs plots. d. AUC curve (red) showing a value of 0.71 when comparing our precited perturbation scores to our experimental validations presented in Fig.6i for n=40 different E-P pairs. e. Scatter plot comparing the predicted perturbation scores and the ABC scores for each of the 40 experimentally tested E-P pairs. Spearman Correlation value of −0.49. Different colors indicate different groups reflecting the concordance or discordance between predictions and experimental validations as shown in Fig.6i. TP: true positive, TN: true negative, FP: false positive, FN: false negative. *Note:* all statistics are provided in Supplementary Table 9.

